# A Single-Nucleus Atlas of Seed-to-Seed Development in Arabidopsis

**DOI:** 10.1101/2023.03.23.533992

**Authors:** Travis A. Lee, Tatsuya Nobori, Natanella Illouz-Eliaz, Jiaying Xu, Bruce Jow, Joseph R. Nery, Joseph R. Ecker

## Abstract

Extensive studies of the reference plant Arabidopsis have enabled a deep understanding of tissues throughout development, yet a census of cell types and states throughout development is lacking. Here, we present a single-nucleus transcriptome atlas of seed-to-seed development employing over 800,000 nuclei, encompassing a diverse set of tissues across ten developmental stages, with spatial transcriptomic validation of the dynamic seed and silique. Cross-organ analyses revealed transcriptional conservation of cell types throughout development and heterogeneity within individual cell types influenced by organ-of-origin and developmental timing, including groups of transcription factors, suggesting gatekeeping by transcription factor activation. This atlas provides a resource for the study of cell type specification throughout the development continuum and a reference for stimulus-response and genetic perturbations at the single-cell resolution.

## Introduction

Multicellular organisms have evolved various organs to perform specific functions required for the organism to survive and flourish. Unlike most animals, plants undergo dynamic post-embryonic organogenesis to form new organs throughout life. Some plant organs are highly specialized and developmental stage-specific, and many organs consist of various cell types with distinct sub-functions. However, some cell types share functionality across diverse organs and developmental stages based on anatomical and physiological features. For instance, within many tissues, epidermal cell types, generally defined as part of the outermost layer of cells of an organ, protect the organ from and interact with external environmental cues. In contrast, internally localized vascular cells are required to transport water and nutrients^1,2^. Yet, we still have a limited understanding of the molecular identities of cell types within the context of many tissues throughout development. The extent of similarity between the transcriptomes of identically classified cells from various tissues and any tissue-of-origin transcriptional signatures remains an open question.

While cell type markers exist for shared cell types present across different tissue types, the form and function of many cell types can differ based on the tissue of origin and the cellular niche from which the individual cells originate. Intra-cell type heterogeneity is exemplified by epidermal cells of the sepals and siliques in Arabidopsis, where epidermal cells of these tissues undergo rapid cellular elongation in scattered populations based on transcript dosages of an epidermal master regulator^3^, regulators of the cell cycle^4^, or in response to fertilization signals^5^, demonstrating that identically classified cells or cell types, can undergo varied developmental programs dependent on internal cues. In addition to molecular information that pertains to cell type classification, cellular states, which can be defined as variations in molecular phenotypes within a cell type that do not impact its developmental potential^6^, are similarly reflected within the transcriptomes of cells. Cell states can be affected by developmental factors, such as cell cycle^7^, tissue/cellular age^8^, the spatial location and cellular neighborhoods within tissues, and external stimuli^9^. Thus, the transcriptional identity of a cell can be defined as the holistic expression of transcripts involved in determining both cell type and cell state. The diversity of cell states across tissues and development remains largely elusive in plants, but the distinction of cell states within individual cell types can be aided by the investigation of diverse tissues, which permits the comparison of commonly classified cell types across tissues and development.

High throughput single-cell RNA-sequencing (RNA-seq) has been demonstrated to provide detailed maps of cell types in plants^10^. Still, its application is currently limited to selected organs, tissues, and cell types^11–15^, with a predominant focus on the Arabidopsis root tip^16–18^, posing a bottleneck toward a comprehensive understanding of cell types and states in this model organism. Characterization of plant cell types and states across organs at the molecular level will be paramount for understanding organ development and function. This motivated us to comprehensively profile cell identities throughout Arabidopsis development, spanning all major organs present over the plant’s entire life cycle (i.e., from seed to next-generation seeds). Here, we present an initial version of a seed-to-seed atlas of Arabidopsis development that may function as a foundational dataset for more focused studies, as well as be further built upon with future single-cell/nucleus transcriptome and multi-omic studies^19^. Our seed-to-seed single-nucleus transcriptome atlas revealed a vast diversity of previously uncharacterized molecular identities that integrate both cell type and cell state, including both universal and tissue-specific cell type marker genes, which will be a powerful platform for hypothesis generation of specific cell populations along the spatiotemporal axis of Arabidopsis development.

## Results

### A comprehensive single-nucleus atlas of the Arabidopsis lifecycle

To generate a comprehensive atlas of Arabidopsis development, we collected six distinct organs that encompass a diverse range of tissues present at several developmental stages and transitions throughout the entire life cycle corresponding to established developmental roadmaps^20–22^, including imbibed and germinating seeds, three-time points of seedling development, developing and fully emerged rosettes, the stem (apical, branched area and stem base), a range of flower tissue (unopened flower buds to fully mature flowers), and siliques (immature to fully elongated green siliques) (Figure 1A). To circumvent the range of plant cell sizes and reduce non-uniform sampling between tissues, we established a universal nuclei extraction protocol, amenable for droplet-based single-nuclei sequencing, which was further optimized for each tissue (see Methods). A total of 801,276 nuclei from the ten organs and developmental time points passed accepted droplet-based single nuclei filtering metrics^23^ and were independently clustered and merged into a global dataset (Figure 1B). A parallel analysis using higher stringency filtering cutoffs resulting in 432,919 nuclei revealed a similar global structure (Figure 1C), demonstrating the overall high quality of the larger dataset but better resolved discrete cell clusters of mixed sample input. Overall, we chose to more deeply sample the seedling tissue, as this sample constitutes an entire organism with a potentially greater diversity of transcriptional states from the presence of above– and below-ground tissues (Figure 1D). Across all tissues and time points, we observed comparable transcriptional complexity between samples (Figure 1E). Simple access to our single-nucleus datasets can be found at our interactive Arabidopsis developmental atlas browser (http://arabidopsisdevatlas.salk.edu/), which enables the exploration of our single-nucleus datasets within internet browsers (Figure 1F).

**Figure 1.**
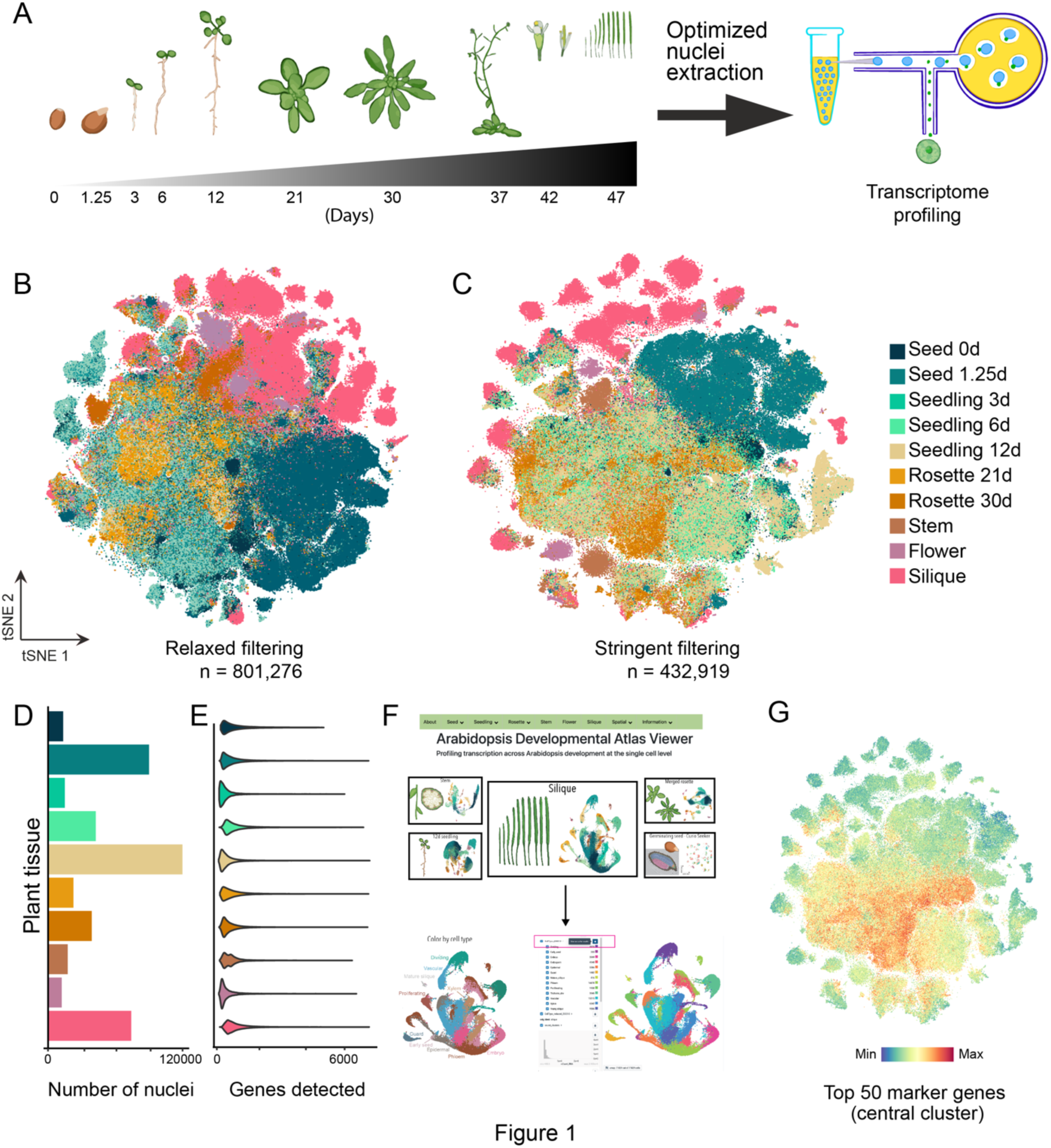
A seed-to-seed atlas of Arabidopsis thaliana. (**A**) Illustration of the collected tissues over the 10 developmental time points spanning the plant life cycle, including: imbibed and germinating seeds (0d and 1.25d), three stages of seedling development (3d, 6d, and 12d), developing and fully expanded rosettes (21d and 30d), the stem (40d) including basal, apical, and branched regions, flower tissue [stages 6-15^21^], and siliques [stages 2-10^20^]. (**B**) relaxed and (**C**) stringent tSNE embeddings of the fully integrated dataset. Nuclei are colored by the dataset of origin. (**D**) Number of nuclei per dataset. (**E**) Violin plot of genes detected per nucleus in each dataset. (**F**) Depiction of web browser access to the Arabidopsis Developmental Atlas Browser and a CELLxGENE instance of the silique dataset. (**G**) Average expression of the top 50 marker genes identified from the large central cluster in the global tSNE.

The global clustering of all ten datasets revealed several distinctly separated clusters, with a large central cluster (Figure 1C). To determine the defining characteristics of the large cluster, we examined the aggregated and individual expression of the top 50 marker genes of these cells (Figure 1G and Figures S1A-H). Many top markers are involved in photosynthesis, suggesting that this biological process profoundly influences the transcriptome of photosynthetically active cells (Figure S1I). Interestingly, the expression of many photosynthesis-related markers within the central cluster had varied expression patterns, suggesting that either photosynthetic function or level of photosynthetic activity contributes to these cells’ transcriptome and clustering. To determine if the expression of the widely expressed photosynthesis-related markers of the central cluster negatively affected the clustering results of the remaining nuclei, we removed the abundantly expressed photosynthesis-related genes and re-clustered the resulting data. However, a large central cluster remained (Figure S1J and K), suggesting a conserved functionality across several populations of individual photosynthetically active cells as demonstrated previously^14^.

### Comprehensive identification of transcriptional identities

To compare cell types across tissues and development, we first independently clustered and annotated cell types within each of the ten datasets, which revealed a total of 183 clusters (Figure 2A). Annotation of individual clusters was performed using the following guidelines. First, we compiled a comprehensive list of marker genes with known cell type or tissue-specific expression patterns in every organ in Arabidopsis (Table S1), including cell type-specific markers recently identified from single-cell RNA-seq studies^16,24–28^. Many of these markers were enriched in specific clusters, which aided cluster annotation (Figure 2B and Figure S2B). We also calculated a cell type enrichment score for each cell type from the list of cell type markers in individual clusters to systematically infer their cell types (see Methods; Figure S2C). Lastly, we manually examined the expression of top cluster markers identified within each dataset to previously generated dissection-based and cell-type-specific transcriptomic studies (TAIR^29^; ePlant^30,31^) to confirm the accuracy of our cluster annotations. Using this approach, we could assign cell identity to many clusters in this large dataset.

**Figure 2.**
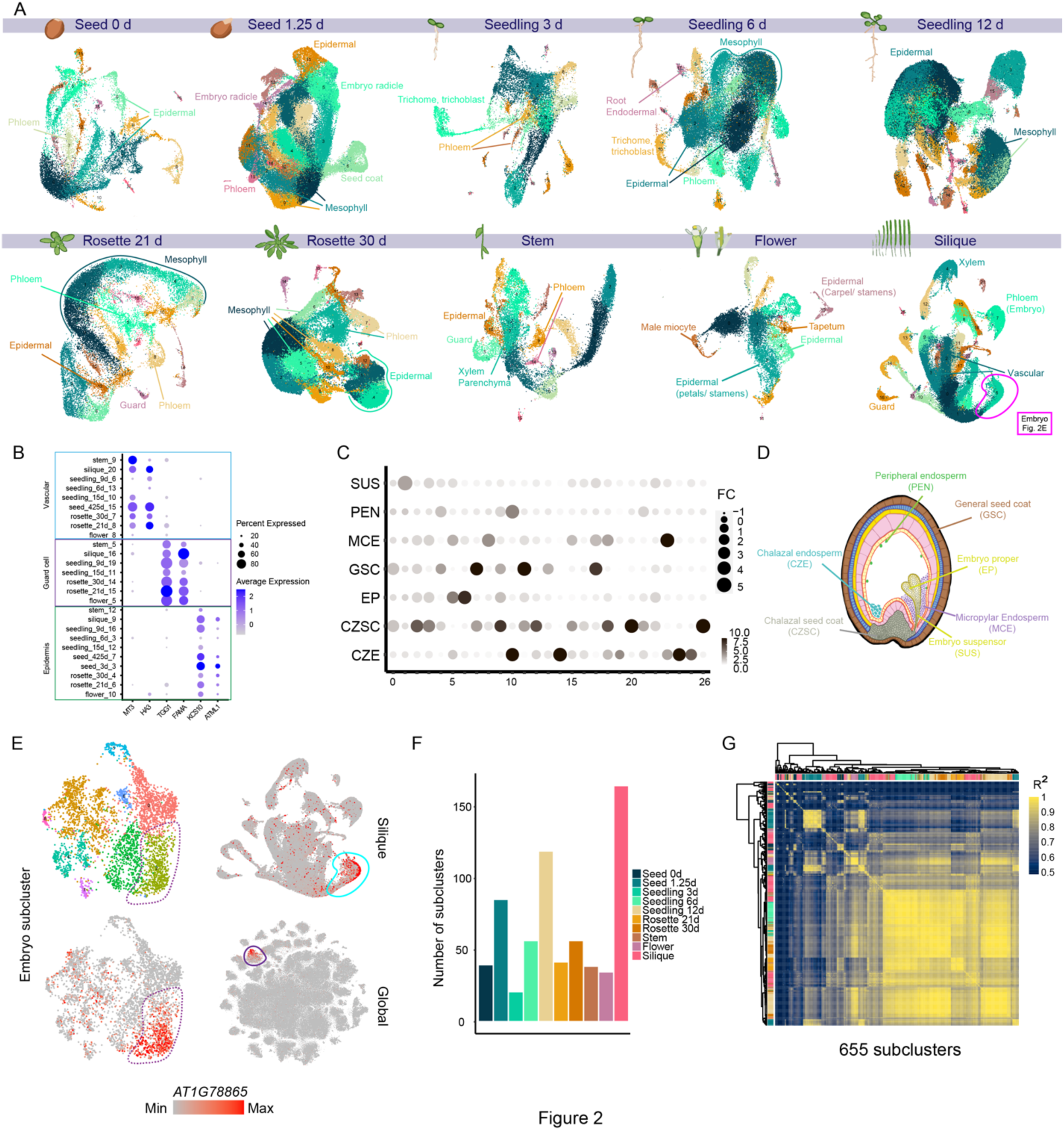
Cellular diversity throughout Arabidopsis development. (**A**) UMAP of each dataset with select high-confidence annotation categories for each dataset. Nuclei are colored by cluster for each dataset. (**B**) Dot plot of known cell type specific markers of vascular cells, guard cells, and epidermal cells, in the corresponding annotated clusters from each tissue. (**C**) Dot plot of literature marker overlap with major clusters identified in the silique dataset. FC, fold change. (**D**) Illustration of early seed development with cell type annotations. (**E**) Subclustering of the annotated embryo cluster from the silique dataset (top left) and the expression of a subcluster marker (*AT1G78865*) within the embryo subcluster (bottom left), silique (top right), and global datasets (bottom left). (**F**) The total number of subclusters identified within each dataset. (**G**) Correlation heatmap of pseudobulk transcriptomes across 655 subclusters. The top color bars indicate the organ of origin.

The organ with the greatest cluster complexity was the silique, where we identified 26 major clusters (Figure 2A and Table S2), which may arise from the diversity of tissues within the fruit organ (fruit flesh and developing embryonic tissue), and from real-time developmental gradients present both across individual siliques and within each silique^32^. Further, within individual siliques, diverse stages of embryonic development from fertilization to zygotic to embryonic development can be present along the longitudinal axis of the fruit based on the timing of fertilization of individual embryo sacs. Using silique cell type-specific marker genes defined by a previous snRNA-seq study^28^, we were able to assign many of the clusters to particular cell types of this organ (Table S2) along with several cell types that comprise the developing embryo and seeds (Figures 2C and 2D).

As the embryo comprises several distinct cell types, we wanted to determine if we could further identify cell types or cellular heterogeneity within this cluster. To further investigate this cluster, we performed a subclustering analysis (subsetting and re-clustering individual clusters, see Methods), which identified nine subclusters, with each subcluster associated with the expression of distinct markers (Figure 2E, left). Examination of a top subcluster marker *AT1G78865*, a noncoding RNA with no predicted function, revealed that expression of this transcript was enriched in only a specific subcluster of the embryo cluster (Figure 2E, bottom left), which may denote an embryo subtype, or reflect a specific developmental stage of an embryo. Interestingly, we found that expression of this embryo subcluster marker was highly enriched, but not entirely exclusive to the embryo cluster in the silique dataset (Figure 2E, top right), suggesting that this noncoding RNA may have additional roles in silique development; yet the expression of this transcript was restricted to a single population of cells in the globally integrated dataset, revealing a highly specific role of this transcript throughout whole-plant development (Figure 2E, bottom right). Together, these results demonstrate that our dataset is sufficient in depth to identify additional transcriptionally unique cell populations through subclustering analyses, and our broad sampling strategy allows for inter-organ comparisons of expression of identified major and subcluster marker genes.

We then systematically subclustered each of the 183 major clusters identified from all organs, resulting in a total of 655 subclusters that may represent both unique cell identities across tissues and development (Figure 2F, Tables S3 and S4). Comparing the correlations of aggregated transcriptomes (pseudo-bulk) of individual subclusters revealed both groups of subclusters, derived from individual organs, that share conserved transcriptional signatures, as well as subcluster groups associated with unique gene expression patterns (Figure 2G). This observation was further supported by analyzing overlapping sets of *de novo* identified cluster marker genes across all 655 subclusters, where we observed particularly high cluster diversity among subclusters of seeds and siliques (Figure S2D-G). Examination of expression patterns of select subcluster markers in the globally integrated dataset confirmed that in many cases, expression was restricted to individual clusters (cell types), but only within subsets of cells of individual clusters, demonstrating the potential for the identification of novel cell populations throughout development (Fig 2E). Together, our seed-to-seed transcriptome atlas captures known cell types and a diverse set of previously uncharacterized cell populations, and enables the investigation of cell type development within and across individual populations of cells.

### Cross-validation and integration of single-nucleus and –protoplast datasets

To assess the quality of our nuclei-based datasets and the accuracy of cluster annotations, we compared our two rosette datasets to a protoplast-based single-cell RNA-seq dataset of 17d-old leaves^25^. Using the protoplast-based dataset as a reference, projection of the nuclei-based datasets and cell type label transfer^33^ revealed that the majority of cell types identified in the protoplast dataset were successfully predicted in the nuclei datasets (Figures 3A and S3A). Of note, equal representation among all clusters between the nuclei– and protoplast-based datasets was not observed (Figure 3B), which may be attributed to differences in the developmental age of the samples (first true leaf vs. whole rosettes), or differences in biases between extraction methods of nuclei and protoplasts, as observed for root tips^34^. Investigation of a protoplast-identified cell type-specific marker (*AT1G16410*, *CYP79F1*) revealed expression in similarly annotated clusters based on cell type prediction in the rosette nuclei datasets (Figure 3C), demonstrating the ability to characterize identical cell types in both protoplast– and nucleus-based datasets. We also determined that the expression of this transcript is restricted to an individual cell type throughout all tissues assayed, demonstrating high levels of cell type specificity throughout development (Figure 3C, bottom right). We were also able to detect markers and co-cluster rare cell populations identified in the protoplast-based study, as evidenced by the abaxial epidermal cells of the proximal midvein, which were identified as a subpopulation of the epidermal cells (Figure 3D). In contrast to the vascular-associated protoplast marker (Figure 3C), expression of the abaxial epidermal marker of the proximal midvein (*AT2G39310*, *JAL22*) was more widely expressed among other tissues in the globally integrated dataset (Figure 3D, bottom right), yet still restricted to cluster specific patterns, suggesting that the expression of this gene is more broadly associated with cellular-populations or – states that are present within other tissues, or stages of development.

**Figure 3.**
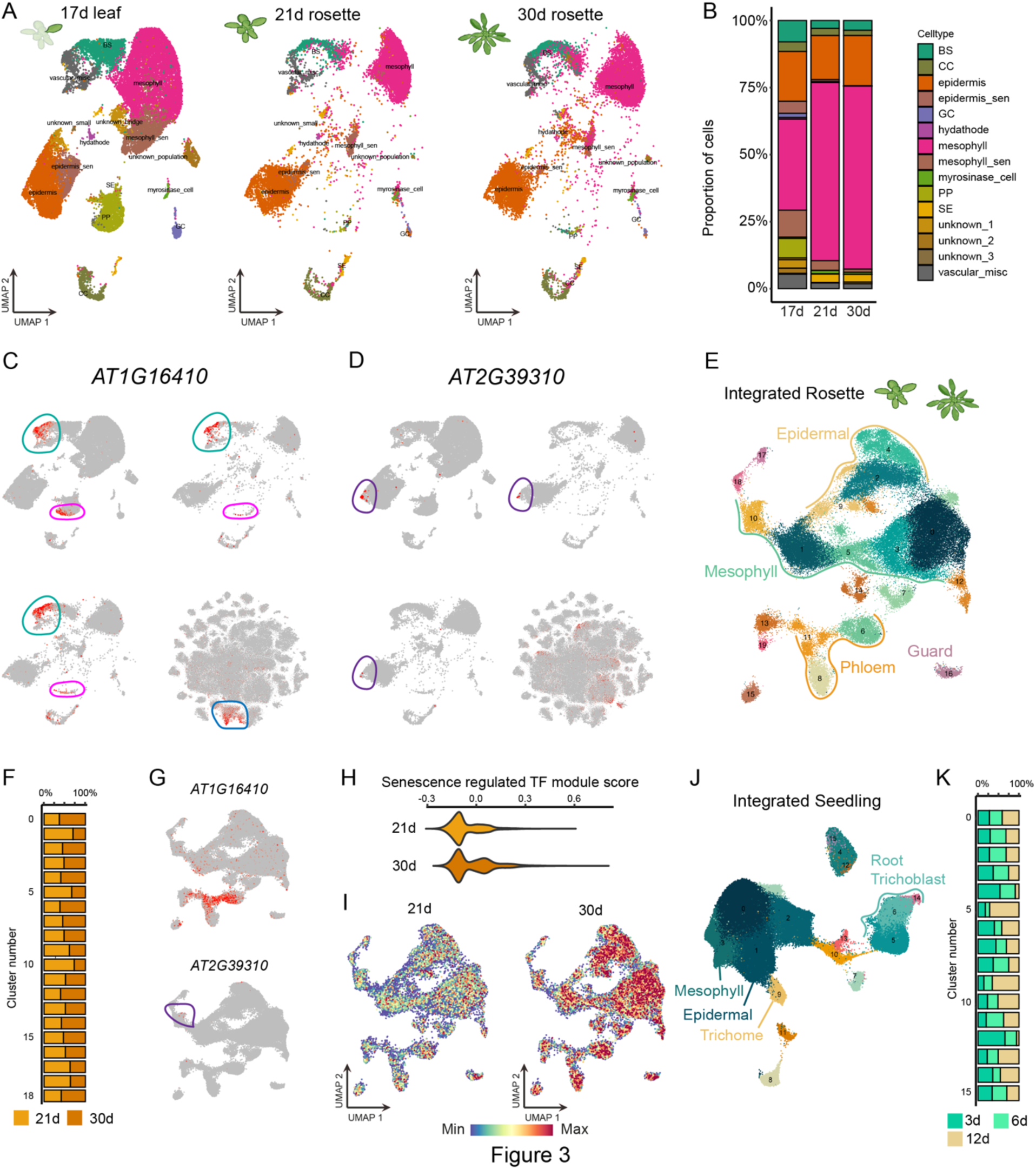
Cross-validation and integration of single-nucleus and –protoplast datasets. (**A**) Integration of a 17d-old leaf single-cell RNA-seq dataset (left) with the 21d– and 30d-old rosette single nucleus datasets (middle and right, respectively). Cells and nuclei are colored by the annotated and validated cell type (single-cell dataset) and the predicted cell type (single-nucleus dataset). (**B**) Percentage of annotated and predicted cell types within each dataset. BS, bundle sheath; CC, phloem companion cell; GC, guard cell; PP, phloem parenchyma; SE, sieve element. (**C** and **D**) Expression of cluster and subcluster markers identified in the leaf single-cell RNA-seq dataset with validated expression in (C) vascular cells (*AT1G16410; CYP79F1*) and (D) abaxial epidermal cells restricted to the proximal midvein (*AT2G39310; JAL22*). (**E-G**) Integration of the 21d– and 30d-old rosette single-nucleus RNA-seq datasets. (F) Distribution of cells per dataset for each cluster. (G) Expression of the vascular and epidermal midvein markers in the integrated rosette dataset. (**H** and **I**) Expression of senescence regulated transcription factors in the 21d– and 30d-old rosette datasets. (H) Violin plot showing the averaged expression of 12 senescence transcription factors^34^ within all nuclei of the 21d– and 30d-old rosette datasets. (I) UMAP of the integrated rosette dataset with the 21d– (left) and 30d-old (right) rosette datasets depicted. Nuclei are colored by the senescence TF expression score in (H). (**J** and **K**) Integration of the 3d-, 6d-, and 12d-old seedling datasets. (J) UMAP of the integrated seedling dataset with select cluster annotations. (K) Distribution of the 3d-, 6d-, and 12d-old seedlings within each cluster.

As several of our datasets constitute developmental time series (seedling and rosette development), we also performed integrative analyses of these samples. Integration of the 21d and 30d rosette samples revealed a number of clusters consistent with the independent clustering of each sample (Figure 3E). All clusters were represented by both datasets (Figures 3F and S3B), and clusters corresponded to previously annotated cell types (Table 2), with expression of the protoplast markers (Figures 3C and D) similarly restricted to cluster and subcluster populations (Figure 3G). After integrating the dataset, our next step was to investigate whether it was possible to discern age-specific variations among these separate developmental time points. Leaf senescence is a highly controlled, multi-step process with known transcriptional regulators that contribute to the irreversible transition from sink to source tissue^35,36^. Examination of transcription factors that positively regulate senescence^35^ within all nuclei from the 21d and 30d rosette samples revealed an overall increase in expression in the older tissue across all nuclei (Figure 3H), demonstrating the ability to identify age-related differences within this integrated dataset. Interestingly, we observed a broad upregulation of these transcripts in the 30d dataset that was agnostic of cluster/cell type, suggesting that the process of leaf senescence may simultaneously affect a majority of cell types present within leaves (Figure 3I) and demonstrating the regulation of a distinct transcriptional and/or cellular state that occurs independent of cell type. Together, integration of our leaf nuclei-based datasets with protoplast-based datasets and across developmental time points has demonstrated that our datasets can directly integrate with protoplast-based single-cell transcriptomic datasets, as well as reveal distinctions across real-time development, which enables the identification of cell type-specific and – agnostic changes in transcription.

### Reconstructing development along real-time and pseudotime

To further explore real-time developmental dynamics of individual cell types, we utilized the integrated seedling dataset (Figures 3J and Figure 4A) that also represents a time series of seedling development (3-, 6-, and 12-day-old seedlings). Integration of the three datasets revealed that all clusters were represented by each sample, demonstrating that the transcriptional identity of major cell types is conserved across these stages of seedling development (Figure 3K, Figure 4A, and Figure S3C). Despite successful integration, the composition of select clusters was skewed by developmental age, including the annotated root hair clusters (Figure 4A). It is well characterized that the timing of root hair (I) specification, (II) emergence, (III) elongation and (IV) maturation is developmentally gated^37^, with root hairs first observable following two days of growth, suggesting that the unequal distribution of the root hair clusters may reflect biological variations in both quantity and degree of root hair differentiation and development. Thus, our time-resolved seedling atlas provided an opportunity to reconstruct this developmental process by trajectory inference and pseudotime analysis, supported by real-time ground truth. Re-clustering of the annotated root hair clusters revealed heterogeneity within this population of cells (Figure 4B). Fitting a trajectory and pseudotime to these cells revealed a continuous gradient of root hair maturation, capturing expression patterns of known root hair developmental stage-specific genes (Figure 4C). Consistent with our prior findings, we observed that nuclei of the youngest seedling dataset (3d) were enriched earlier in pseudotime. In contrast, nuclei of the older seedlings progressively populated later intervals of pseudotime (6d and 12d) and at greater proportions (Figure 4D). Clustering of genes differentially expressed over pseudotime revealed three expression patterns, with groups of genes (gene modules) that largely correlated to these three stages of root hair development (Figure 4E and Table S5). Of note, the *de novo* identified gene groups corresponding to psuedotime intervals are not enriched for processes related to root hair development, but rather broader GO terms such as anatomical structure process and glycerolipid metabolic process, implicating novel roles of those genes in root hair development (Figure S4A-C).

**Figure 4.**
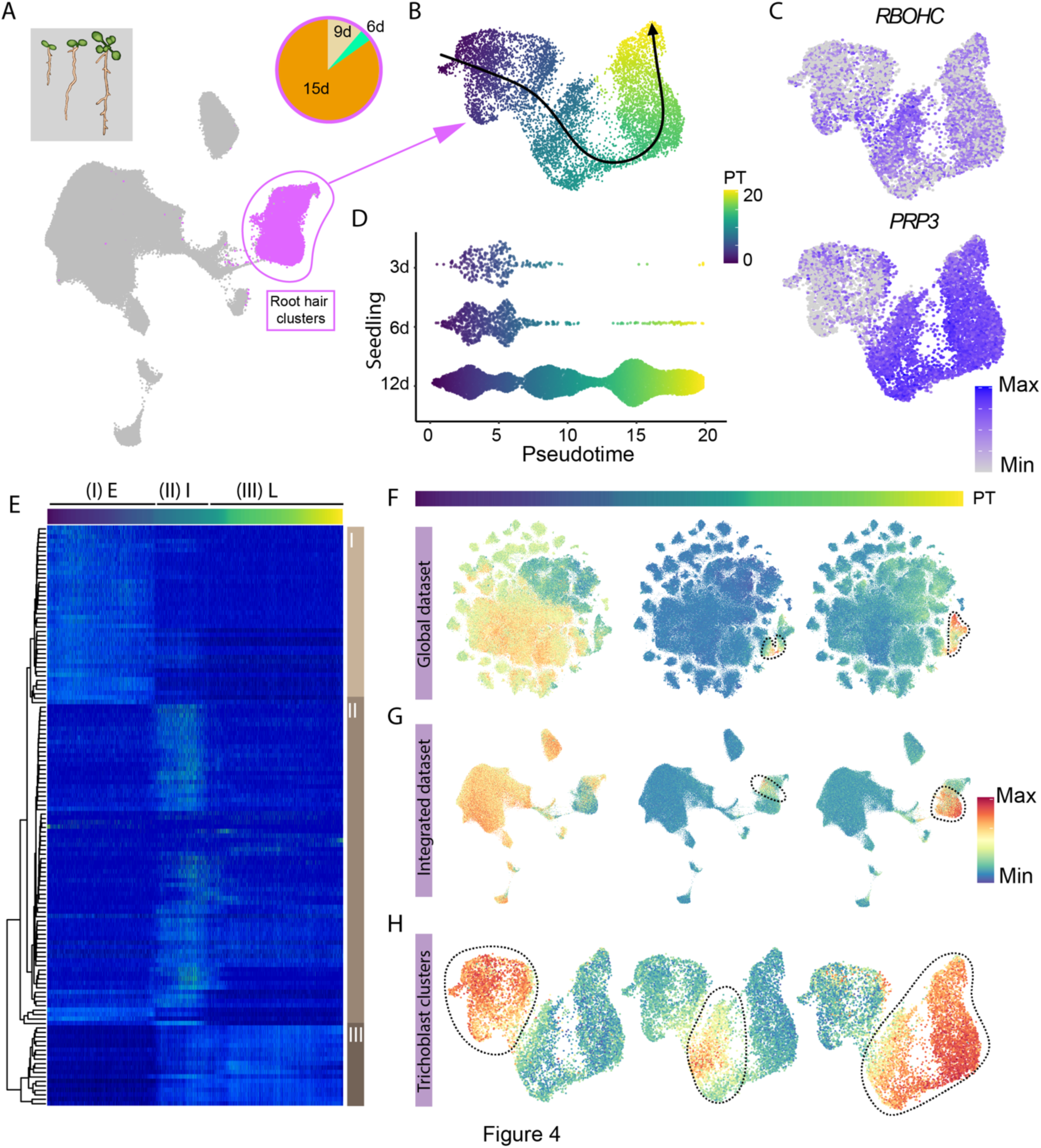
Transcriptional programs of root hair development across the entirety of Arabidopsis development. (**A**) Integrated clustering of the 3d, 6d, and 12d seedling datasets. Root hair (trichoblast) clusters are circled, and the distribution of nuclei within the trichoblast clusters among developmental time points is depicted. (**B**) Re-clustering and pseudotime trajectory of annotated root hair clusters. (**C**) Expression of known marker genes of root hair development (*ROBHC* [*AT5G51060*] and *PRP3* [*AT3G62680*]). (**D**) Distribution of root hair cells from 3d-, 6d-, and 12d-old seedlings across pseudotime. (**E**) Heatmap of 170 genes differentially expressed along pseudotime of root hair cells. Three major classes of genes corresponding to (I) E=early, (II) I=intermediate, and (III) L=late pseudotime, are depicted. (**F**-**H**) Expression of the three gene modules identified from pseudotime in the (F) global, (G) integrated seedling, and (H) root hair re-clustering.

Next, we sought to determine if these spatiotemporally regulated gene modules have conserved expression in other tissues and/or cell types. Mapping the average expression of each gene module to the integrated seedling and global datasets revealed patterns of both highly specific and broad roles of these genes across development (Figures 4F-H). For example, the gene module corresponding to intermediate root hair development (intermediate; II) was exclusively expressed within this small population of developing root hair cells throughout all tissues and developmental time points assayed (Figures 4F-H; top middle, circled), suggesting that the expression of these genes may be unique to root hair cells throughout the entirety of Arabidopsis development in plants grown in controlled conditions. Conversely, genes corresponding to both early and late stages of root hair development were more broadly expressed in other tissues and cell types (Figures 4F-H, left and right panels), suggesting that the expression of these genes are more broadly utilized throughout development. Overall, by inclusion of broad tissues and developmental stages, we were able to identify groups of genes with shared function across cell types of various tissues, in the case of root hair initiation and late-stage development. Conversely, we were also able to identify groups of genes that are uniquely utilized by a single cell type at discrete stages of development, in the case of intermediate root hair development, demonstrating that the expression of some genes may be under the combined control of cell type-specific and developmental timing-specific regulation.

### Cell type– and tissue-of-origin-specific genes define the transcriptional identities of cells

It is known that various organs may consist of recurrent cell types with shared functions and morphology (e.g., guard cells). However, the extent of similarity between the transcriptomes of commonly shared cell classes from distinct tissues is currently unknown. To investigate this question, we queried the globally integrated dataset (Figure 1C) that enables the identification of commonly shared cell types originating from varied tissues throughout development. We observed shared strong transcriptional identities across organs in some cell types, such as phloem companion cells and guard cells, characterized by expression of the marker genes *SUC2* (*AT1G22710*^38,39^) and *FAMA* (*AT3G24140*^40^), respectively (Figure 5A). Independent subclustering of both the companion cell and guard cell populations revealed similar results, with the seedling and rosette tissues generally co-clustering within each developmental time series (circled), while nuclei isolated from other tissues, exemplified by the silique tissue, clustered separately (Figures 5B and C), suggesting tissue-specific transcriptional influences within individual cell types. The consistent segregation of cells from the silique tissue in the companion cell, guard cell, and other discrete clusters of mixed sample identity (Figures S5A-C) motivated us to question what kind of genes–whether tissue-specific housekeeping genes or unique subsets of genes within each cell type–contributed to the intra-cell type heterogeneity between cells of various tissues.

**Figure 5.**
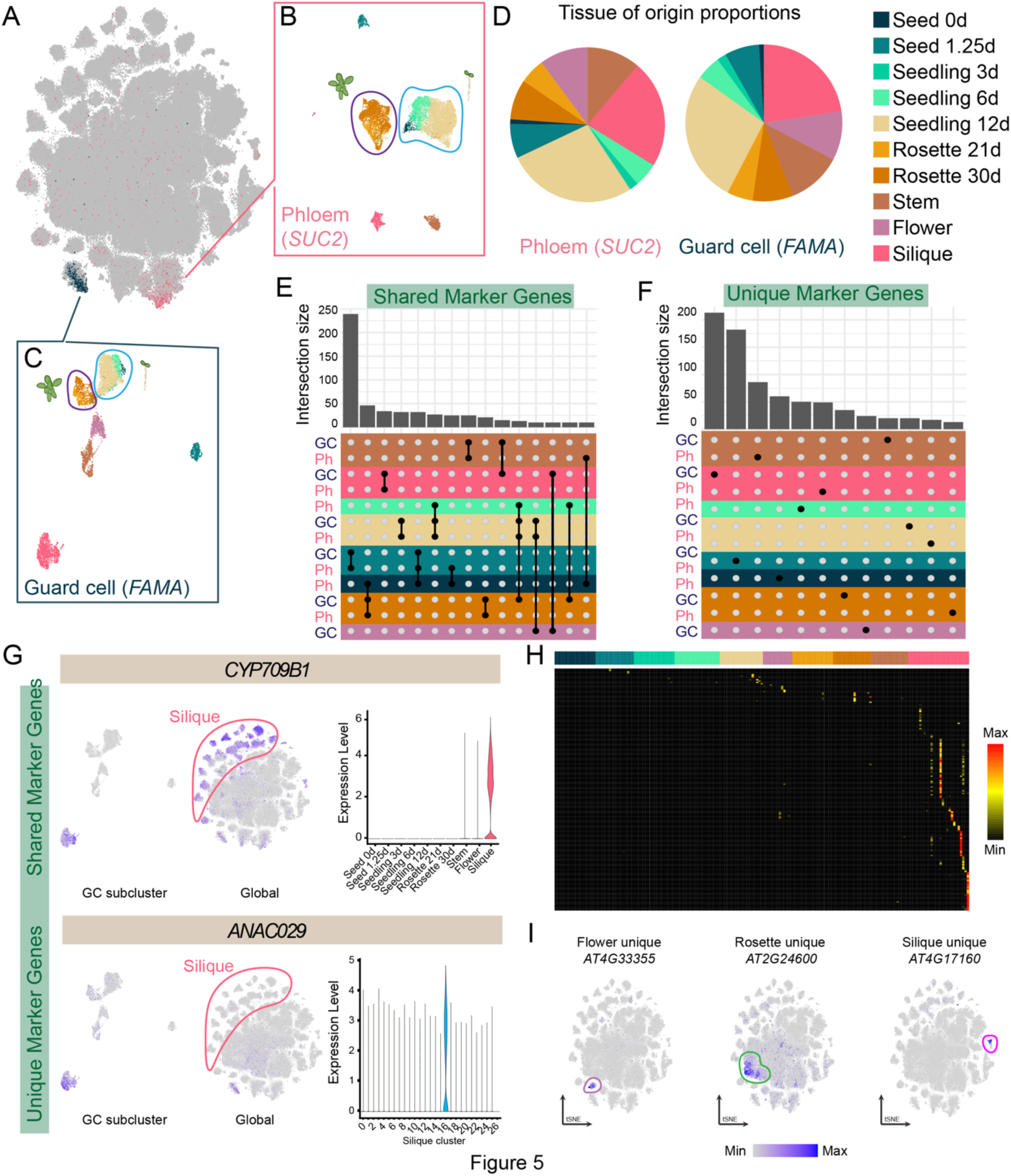
Identification of tissue– and cell type-specific transcriptional signature. (**A**) tSNE of the global dataset with expression levels of the stomatal lineage marker *FAMA* (blue, [*AT3G24140*]) and phloem marker *SUC2* (pink, [*AT1G22710*]). (**B** and **C**) Re-clustering of the (B) phloem and (C) guard cell lineage clusters. Cells are colored by tissue of origin. (**D**) Distribution of cells per dataset in the phloem and guard cell lineage clusters. (**E**) Quantification of subcluster markers shared between phloem and guard cell lineage subclusters. Groups of subclusters with greater than ten overlapping subcluster markers are depicted. Black dots indicate the overlapping group(s) of subclusters. Color indicates the dataset of origin with the highest proportion of cells for each subcluster. (**F**) Quantification of markers uniquely identified as a subcluster marker in only one subcluster of phloem and guard lineage cells. Subclusters with at least ten uniquely identified genes are depicted. (**G**) Expression of a representative shared tissue (*CYP709B1* [*AT2G46960*]) and unique marker (*ANAC029* [*AT1G69490*]) in the stomatal lineage subclustering, global integrated dataset, and expression within each tissue dataset. (**H**) Heatmap showing expression of genes uniquely enriched within only one of 183 clusters. (**I**) Expression of unique cluster markers from the flower (left), rosette (middle), and silique (right) datasets at the global clustering level.

Comparing the expression of *de novo* identified subcluster markers of companion and guard cells amongst all subclusters of these two cell types revealed two classes of genes; (I) shared – markers identified in two or more subclusters (Figure 5E and Table S6) and (II) unique – markers identified in only one subcluster (Figure 5F and Table S6). For example, *CYP709B1* (Figure 5G, top) and *ANAC029* (Figure 5G, bottom) were uniquely identified as subcluster markers of silique guard cells (Figure 5G, left), but in the context of the whole-plant development, *CYP709B1* was expressed more broadly, yet only within cells of the silique tissue (“shared marker gene”), while *ANAC029* expression was restricted to only guard cells of the silique within the context of whole-plant development (“unique marker gene”) (Figure 5G, middle). Consistent with the silique example, “shared marker genes’’ of other tissues were also associated with tissue-specific enrichment at the pseudobulk level (Figure S5F), demonstrating that tissue-of-origin signatures can influence transcriptional identity within individual cell types (Figure S5C). These results indicate that for some cell types, as demonstrated with phloem companion and stomatal lineage cells, the transcriptional identity is defined by either cell type– or tissue-of-origin marker genes, or the intersection of them. An extension of this analysis to include additional clusters (ten clusters, Figures S5A-C) yielded similar results, identifying both tissue-shared transcripts, as well as transcripts uniquely expressed within only one cell type at a single time point of development (Figures S5D-F and Table S6).

To systematically identify markers that are unique to a single tissue type, we compared the pseudo-bulk expression of marker genes for each of 183 major clusters (Figure 5H, see Methods). Out of 4,528 genes that were identified as markers of at least one of the major clusters, 331 (7.3%) genes were uniquely expressed in a single tissue type (Table S6). Consistent with the unique expression patterns in the pseudo-bulk data, plotting the expression of these markers in the globally integrated dataset revealed that the expression of many of these markers was restricted to individual clusters and/or subsets of clusters, demonstrating that our atlas can identify transcripts that are uniquely expressed in specific cell types at specific developmental time points (Figure 5I). Together, we revealed classes of genes uniquely expressed within a single cell type of a single tissue, as well as tissue-of-origin transcriptional signatures that holistically define the transcriptional identity of cells.

Of note, we observed the largest quantity of shared subcluster markers between cell types of the germinating seed (1.25d) (Table S6). A deeper investigation of the shared markers of germinating seeds revealed that many of the identified genes with shared expression across cell types encode a diverse range of ribosomal protein subunits. The unified expression of ribosomal proteins across cell types in the germinating seed dataset coincides with global increases in ribosome biogenesis and translation during seed germination in both Arabidopsis^41,42^ and Barley grains^43^. While the expression of many ribosomal protein genes is present across several cell types of germinating seeds, this was not observed for all of the other tissues assayed, where subsets of ribosomal proteins were expressed only in specific cell types among all tissue and organs assayed, supporting the hypothesis that the heterogeneous composition of ribosomes is developmentally regulated^44^, and may explain the higher levels of homogeneity within this tissue. As seed development progresses from imbibition to germination, embryo cell types progress through reprogramming^42^, suggesting that a snapshot of cellular state was captured in our germinating seedling dataset.

### Transcription factor specificity across development

We systematically analyzed TF expression across the identified cell populations to understand cell type and organ-specific gene regulatory mechanisms. Many TFs showed organ-specific expression (Figures 6A and B), but we also observed heterogeneity within organs (Figures S6A-C) and among major clusters (Table S7). For instance, *AT3G15510* (*ANAC056*) was expressed specifically and globally in the silique tissue (Figure 6C); *AT3G62340* (*WRKY68*) was also specifically expressed in the silique, within a subset of cells, but also the entire context of development (Figure 6D). Notably, we identified TFs expressed in specific clusters of each organ. TFs such as *FAMA*, *WRKY23*, and *BIM1* were highly expressed in clusters mostly annotated as guard cells in seven different samples (Figure 6E; highlighted in red), suggesting that some TFs have conserved cell type-specific roles across various organs and development.

**Figure 6.**
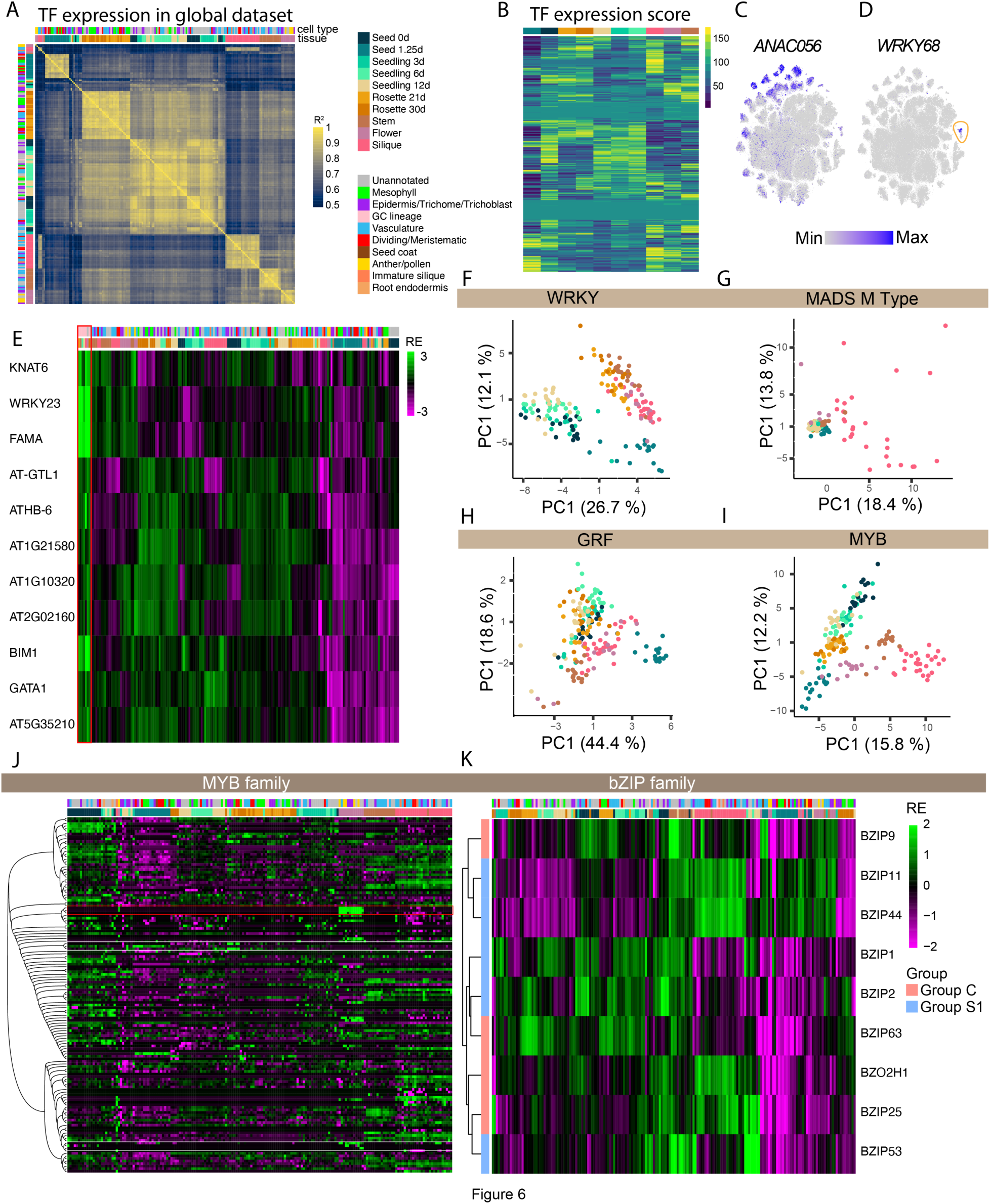
Diverse expression patterns of TFs across cell types and organs. (**A**) Heatmap showing pair-wise correlation of pseudo-bulk TF expression between major clusters of individual samples. Pearson’s correlation coefficient values (R^2^) are shown. (**B**) Averaged ranks of TF expression in the clusters derived from each tissue of origin. A higher value indicates more global and dominant expression of a TF in a tissue of origin. (**C** and **D**) Expression of TFs, *AT3G15510* (*ANAC056*) and *AT3G62340* (*WRKY68*), that are specifically expressed in the silique visualized in a globally clustered tSNE. A cluster with specific gene expression was circled in D. (**E**) Heatmap showing pseudo-bulk expression of marker TFs of global cluster 21 (guard cell cluster; Figure 5C). Clusters that showed strong expression of these markers were highlighted in red. (**F**-**I**) Principal component analysis of TF family expression in each cluster. Each dot represents a major cluster colored by a tissue of origin. TF family names are indicated in the top bars. (**J**) Heatmap showing expression of MYB family TFs in all major clusters. Genes were ordered by the phylogenetic relationships defined by their amino acid sequences. An example of genes that showed tissue specific expression were highlighted in red. Examples of closely related genes that showed contrasting expression patterns were shown in white. (**K**) Heatmap showing relative expression of bZIP family TFs that belong to Group C and S1. For each heatmap, top bar indicates tissue of origin and/or cell type annotation. (E, J, K) Expression was shown as log_2_ relative expression (RE).

We next asked whether TFs within the same TF family show organ specificity and within-organ heterogeneity. We performed principal component analysis on the expression of the members of 70 TF families in individual clusters. We found both cases where TF family members showed either shared or diverse cell type/organ specificity (Figures 6F-I and Figure S6D). For instance, expression patterns of WRKY TFs are primarily separated into three organ groups (regardless of cell type): (I) germinating seeds (1.25d), (II) imbibed seeds (0d) and seedlings, and (III) rosette, stem, flower, and silique (Figure 6F). Cell type/state heterogeneity was observed within each organ group (Figure 6F). Some TF families have largely expanded and showed high degrees of heterogeneity (Figures 6J and K and Figure S11E and F). We asked whether sequence similarities between TFs can explain their organ/cell type specificity. We constructed a phylogenetic tree for each TF family based on amino acid sequence and overlaid gene expression. Interestingly, we found cases where phylogenetic relationships of genes within TF families can explain their organ/cell type specificity, as clusters associated with cell types and organ formed grouped when TF family members were clustered phylogenetically (Figure 6J; highlighted with red boxes). Conversely, we found cases where closely related TFs showed contrasting expression patterns (Figure 6J; highlighted with white boxes). These results imply the varying history of the evolution of tissue specificity among TFs.

It is known that some TFs activity depends on interacting partners for function. For instance, basic leucine zipper (bZIP) TFs recognize target DNA sequences by forming various homo– and hetero-dimers^45^. A previous study showed that Group C bZIP TFs (bZIP9, bZIP10, bZIP25, bZIP63) do not bind DNA as homodimers but form heterodimers with Group S1 members (bZIP1, bZIP2, bZIP11, bZIP44, bZIP53) to bind unique DNA sequences^46^. We found that genes in each group showed varying cell type/tissue specificity (Figure 6K). Interestingly, Group C TFs were clustered with at least one Group S TF with overlapping expression patterns (Figure 6K), implying the co-evolution of cell type/organ specificity in these dimerization pairs. Together, our dataset will be valuable for analyzing the neo/sub-functionalization of TFs over the course of gene family expansion, which has previously been suggested for some genes^47^.

### Spatial mapping of cell types

Annotating cell types and states to specific clusters identified in snRNA-seq requires *a priori* knowledge of previously validated cell type-specific markers by imaging– and dissection-based analyses. However, such ground truth cell type markers are unavailable for many organs and cell types within the entire Arabidopsis lifecycle. Therefore, to validate our cluster annotations, we utilized two spatial transcriptomics technologies to simultaneously validate cell type-specific expression patterns of several *de novo* identified clusters from two datasets associated with highly dynamic stages of growth: germinating seeds (1.25d seeds) and siliques.

To spatially profile the transcriptome of germinating seeds (1.25d) (Figure 2A), we utilized a sequencing-based spatial transcriptomics platform with 10 µm spatial resolution to profile cell type and cell layer-specific transcriptomes^48^. Plotting the transcript detection of spots in spatial coordinates revealed shapes resembling seed and embryo structures (Figures S7A and B). Dimensionality reduction and *de novo* clustering of the filtered spatial transcriptomics data revealed two major groupings of clusters (Figure 7A and Table S2). Spatial mapping revealed that these clusters broadly correspond to the cotyledons (clusters 0 and 5), root tip region (cluster 1), the epidermis (cluster 2), seed coat (cluster 3), and the provasculature (cluster 4) (Figures 7B-D). Mapping the expression of cluster markers corresponding to the cotyledon and epidermal clusters (clusters 0 and 2; Figure S7) onto the matched droplet-based single-nuclei dataset similarly revealed cluster-specific expression patterns to the correspondingly annotated cell types (Figure 7E), demonstrating the ability to annotate the droplet-based clusters accurately.

**Figure 7.**
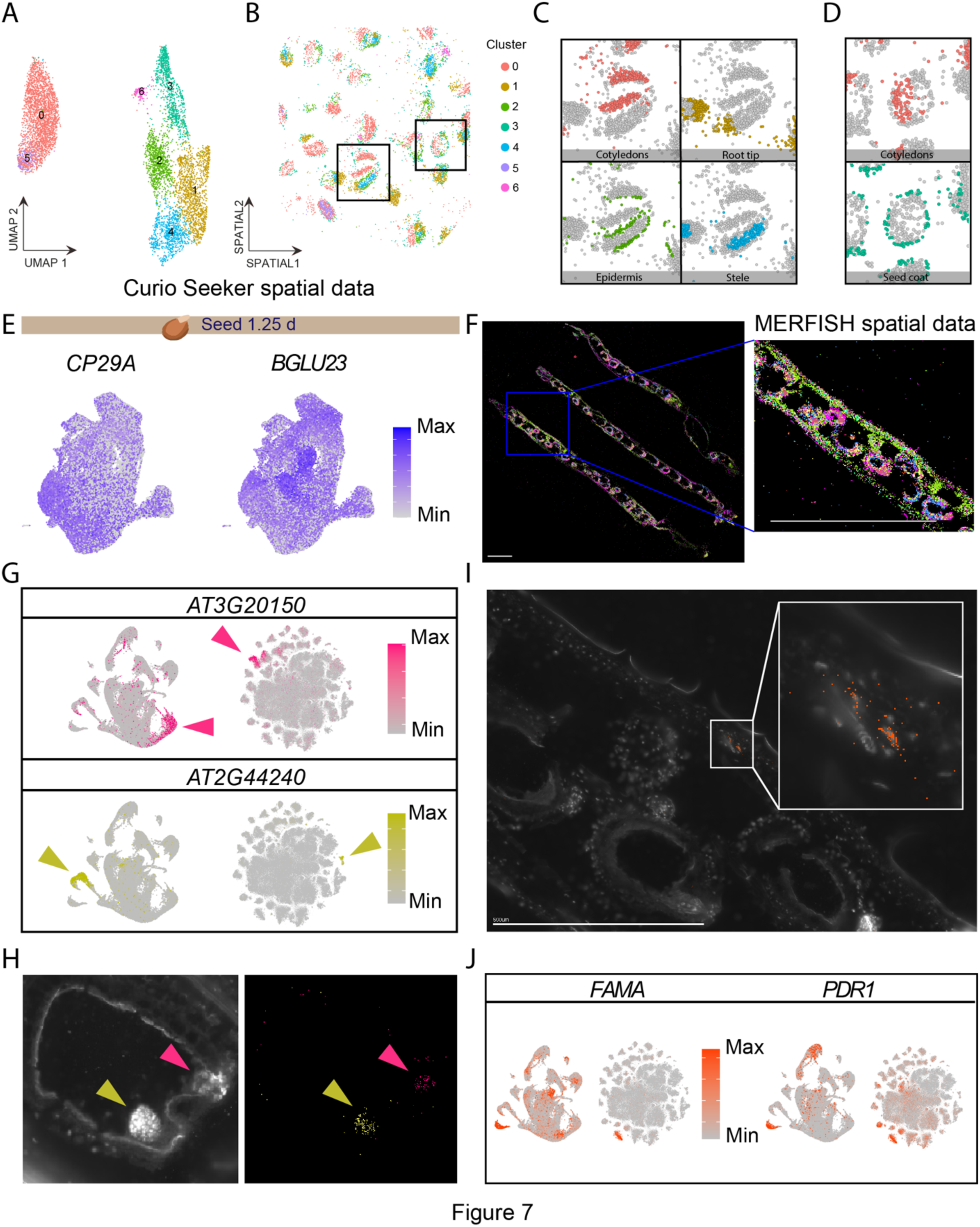
Spatial transcriptomics annotates silique cell types and validates de novo marker genes. (**A**) Dimensionality reduction and *de novo* clustering of spatial transcriptomics data of 1.25-day-old germinating seeds. (**B**-**D**) Mapping of clusters to spatial coordinates with a focus on clusters corresponding to the (C) embryonic seedling and (D) seed coat. (**E**) Expression of *de novo* identified spatial cluster markers corresponding to cotyledons (left, *CP29A* [*AT3G53460*]) in the matched single-nucleus droplet-based dataset (Figure 2A). (**F**) Visualization of all 140 transcripts targeted with MERFISH in siliques. Scale bar = 1 mm. (**G**) Expression of *de novo* identified marker genes of silique clusters predicted to be the embryo (*AT3G20150*) and chalazal endosperm (*AT2G44240*) in the (left) silique and (right) globally clustered datasets. (**H**) (Left) DAPI staining of an embryo within the silique. (Right) Spatial mapping of predicted marker genes of embryo (yellow) and chalazal endosperm (pink). Scale bars = 0.2 mm. (**I**) Spatial mapping of identified guard cell marker genes. The area in the white box was enlarged. Scale bar = 0.5 mm. (**J**) Expression of two *de novo* identified guard cell lineage markers shown in the silique dataset (left) and the global dataset (right).

As fully elongated siliques simultaneously represent dynamic growth patterns associated with both small (egg sac; embryo) and large tissues (∼15 mm silique when fully elongated), we opted for an imaging-based spatial transcriptomics method (MERFISH; see Methods) that can accommodate higher resolution of small cells as well as larger tissue areas to simultaneously visualize the expression of many transcripts on a tissue section at single-molecule resolution. We selected 140 *de novo* identified silique cluster marker genes and spatially mapped them on 10 µm tissue sections of fully expanded green siliques (Figure 7F; Table S1). Marker genes of clusters predicted to be embryo and chalazal endosperm (Figure 2C-D, Table S2) were successfully mapped in expected regions of the tissues (Figures 7G and H). Among these markers, *AT2G44240* has been validated as a chalazal marker by *in situ* hybridization^28^. We also identified cells in the stomatal lineage based on the expression of the guard cell marker gene *FAMA*^40^ and validated the *de novo* identified guard cell marker gene *AT3G16340* (*PDR1*) (Figures 7I and J). These results validate our cell type prediction and, *de novo* identified marker genes of these cell types. Mapping the expression patterns of these highly specific genes revealed that many of these cell type-specific transcripts of the silique are uniquely expressed in this tissue throughout all stages of Arabidopsis development assayed in this study, highlighting the ability to investigate cell type and developmental-specific expression patterns in our dataset (Figures 7G and J). In summary, spatial transcriptomics validated *de novo-identified* cluster markers and facilitated cluster annotation in these complex organs.

## Discussion

In this study, utilizing single-nucleus sequencing technologies, we sought to characterize cell identities across a diverse range of tissues and organs along the entire life cycle of the model plant Arabidopsis, with a focus on capturing and comparing diverse cell populations across organs and developmental stages, as opposed to creating fully-annotated atlases of individual organs. With a broad sampling strategy, our single-nucleus atlas of Arabidopsis development provides a resource of wide use to the Arabidopsis and plant biology community for hypothesis generation and reference for future single-cell genomics studies. Successful integration of our data with published single-cell protoplast-based datasets suggests that our atlas could be used as a reference data onto which future single-cell omics data can be added for more thoroughly annotating cell types and investigating cell type-specific responses to environmental stimuli.

Across the diverse tissues and developmental time points assayed, we were able to identify not only known cell types based on well-characterized marker gene expression but also a wealth of novel transcriptional identities. 655 cell populations identified across ten distinct tissue types showed diverse transcriptome patterns. For each of these 655 individual cell populations, we identified *de novo* marker genes, which can be used for future characterization of the cell populations.

Our developmental atlas also includes discrete time points of seedling and rosette development, which enables the investigation of these tissues through real-time development. Integration of both datasets revealed clusters corresponding to known cell types, which enabled the identification of transcriptional signatures that were highly unique to specific subsets of cells throughout the entirety of the Arabidopsis lifecycle, in the case of root hair development. Using the integrated rosette dataset, we also identified age-related differences in the expression of known regulators of leaf senescence that are regulated independently of cell type. These results suggest that the process of cellular senescence confers an additional layer of transcriptional identity and is potentially indicative of a distinct cellular state within aging leaves. Together, our time series datasets provide a foundation for the integration of a variety of developmental time points for seedling and leaf tissues and enables the investigation of age-related distinctions within individual cell types.

Comparative transcriptome analyses of common cell classes across tissues is a unique analysis enabled by our comprehensive atlas, which revealed that cell identities can be holistically defined by both cell type and developmental or tissue-of-origin gene expression signatures. The presence of tissue-of-origin-specific transcripts may indicate high levels of coordinated activity between cells within tissues, such as cellular elongation within developing siliques, and may serve in the designation of discrete cellular states.

Conversely, transcripts uniquely expressed in only one cell type/state of a single organ are of particular interest as they shed light on cell identity-specific functions and gene regulatory mechanisms, which can be instrumental for targeted precise engineering of specific cell populations to minimize pleiotropic effects of plant genetic engineering. Interestingly, cell types of the terminal reproductive tissues were associated with greater quantities of uniquely expressed transcripts compared to other tissues. It is possible that the high level of uniquely expressed transcripts may be due to the presence of diverse cell types and cell populations within these organs, as these organs constitute both germ– and non-germ-line cells. Further, reproductive tissues are associated with the production of diverse and unique secondary metabolites^49,50^ throughout Arabidopsis development, and may in part be a basis for the observed unique expression patterns. Within the groups of unique marker genes were many TFs, some of which showed phylogenetic conservation, which may explain their organ specificity. Further phylogenetic analysis of large gene families could reveal evolutionary mechanisms underlying the diversification and specification of gene expression across cell types and organs.

Despite rigorous efforts to annotate the identified clusters, some clusters in our atlas still lack definitive cell type or state identities, which is the case in large-scale atlas projects of other organisms. For instance, in the fly cell atlas, >70% of cells of a certain tissue type remain unannotated^51^. Spatial analysis of cluster marker genes offers a potent approach for better understanding these unidentified clusters. The application of two spatial transcriptomics technologies to the germinating seed and silique tissues revealed that high-resolution spatial studies can disentangle plant organ heterogeneity, as evidenced by the validation of both cluster markers and cluster annotations in these highly dynamic tissues. As *in situ* spatial transcriptomic methodologies require knowledge of up to thousands of transcripts with distinct expression patterns, our datasets also function as a foundation for future spatial transcriptomics experiments. Methods such as MERFISH and PHYTOMap^52^, which can spatially map dozens of genes in thicker tissues in 3D, will place diverse cell populations identified in the present study into the native tissue context. Overall, our dataset highlights the complexity of cell types throughout the entirety of the development of an organism. Our dataset also serves as a foundation for future studies that seek to characterize these tissues and developmental stages further, or investigate stress or stimulus-driven responses, with access to the processed datasets for rapid exploration in a web-based interface available at our web portal (http://arabidopsisdevatlas.salk.edu/).

## Acknowledgments

We thank Genevieve Zzyzyx for the illustrations. Thanks to James Walker for annotating pollen cells and improving the manuscript and to Hanqing Liu for reviewing and providing input. T.N. was supported by the Human Frontiers Science Program (HFSP) Long-term Fellowship (LT000661/2020-L). N.I-E is a research fellow at the George E. Hewitt Foundation for Medical Research and an Awardee of the Weizmann Institute of Science – Israel National Postdoctoral Award Program for Advancing Women in Science. J.R.E. is an Investigator of the Howard Hughes Medical Institute.

## Author contributions

T.L., T.N., and J.R.E. conceived and designed the study and experiments. T.L. and T.N. performed method development and data acquisition with technical support by B.J. and J.R.N. T.L., T.N., N.I.E, and J.X. analyzed data. T.L., T.N., N.I.E., and J.R.E. wrote the manuscript.

## Data and materials availability

All datasets are available for browsing at arabidopsisdevatlas.salk.edu. Raw and processed datasets are available at GEO (accession number GSE226097).

## Supplementary Materials

### Materials and Methods

Figure S1 Characteristics of the central cluster.

Figure S2 Clustering of individual datasets.

Figure S3 Projection and integration of time series datasets.

Figure S4 GO terms of root hair pseudotime expression modules.

Figure S5 Cross-tissue cell type analysis of ten cell type populations.

Figure S6 Transcription factor (TF) analysis.

Figure S7 Spatial transcriptomics of the 1.25d germinating seed.

## Supplemental Tables

Table S1. Cell type and tissue marker genes collected from the literature and this study

Table S2. Table of cluster markers and cell type annotation of individual tissue datasets

Table S3. Pseudobulk expression values of 183 major clusters

Table S4. Pseudobulk expression values of 655 subclusters

Table S5. Genes differentially expressed over pseudotime of root hair development

Table S6. Table of uniquely expressed and shared subcluster markers identified from the cross-cell type analysis

Table S7. List of transcription factors identified as cluster markers from the 183 major clusters

## The PDF file includes

Materials and Methods

Supplementary Text

Figures S1 to S7

References

## Other Supplementary Materials for this manuscript include the following

Tables S1 to S7

## Materials and Methods

### Plant growth and sampling

For the imbibed seeds (0-day-old), germinating seeds, (1.25-day-old), and 3-, 6-, and 12-day-old-seedling developmental time points, Arabidopsis Col-0 seeds (CS7000) were surface sterilized by incubation in 70% (v/v) ethanol for 5 min, followed by incubation in 50% (v/v) bleach and 0.01% (v/v) TWEEN-20 for 5 min, followed by triplicate washes in double distilled water for 5 min. Sterilized seeds were placed onto 1 X Linsmaier & Skoog media (1.0 LS salts [Caisson Labs], 0.8% agar [w/v], 1% w/v Suc., pH5.7) in 120 mm^2^ petri dishes at a density of 100 seeds per row for 0d and 1.25d samples, and 30 seeds per row for 3d, 6d, and 12d samples. The imbibed seeds were stratified by incubation at 4°C for 3 d in complete darkness. Following stratification, the plates were placed in a long day (16h light/8h dark) growth chamber at 23°C. For the rosette, flower, and silique tissue, seeds were sterilized identically to the seed and seedling samples, and were directly sown onto soil and were grown at 23°C in long-day conditions.

### Nuclei extraction and single-nucleus sequencing

Nuclei purification buffer (NPB; 20mM MOPS (pH 7), 40mM NaCl, 90mM KCl, 2mM EDTA, 0.5mM EGTA, 0.5mM spermidine, 0.2mM spermine, 1:100 protease inhibitor cocktail, 2% BSA, and 1:1000 SUPERase IN) was prepared fresh and chilled on ice. All the following procedures were performed on ice or at 4°C. Tissues were chopped in 500-1,000 µL of cold NPB with a razor blade on ice for 5 min to release nuclei and incubated in 10 mL NPB. For the 0d and 1.25d seed samples, the nuclei were extracted in 5 mL NPB with a Dounce homogenizer with 20 strokes of the loose pestle followed by 20 strokes of the tight pestle. The crude nuclei extract was sequentially filtered through 70 µm and 30 µm cell strainers. BSA, Triton X-100, and NP40 were added to final concentrations of 2%, 0.05%, and 0.05%, respectively, and the extract was incubated at 4°C for 10 min with rotation. Nuclei were pelleted by centrifugation at 700 x g for 5 min with a swing-rotor centrifuge, The pellet was resuspended in 10 mL of NPBd (NPB, 0.05% Triton X-100, 0.05% NP40, 2% BSA) by pipetting and incubated for 10 min, followed by centrifugation at 700 x g for 5 min. For seedlings and rosettes, additional washes in NPBd was performed until the pellet became translucent. The pellet was then resuspended in 1 mL NPB and centrifuged at 50 x g for 3 min in a fixed-rotor centrifuge to pellet non-nuclei debris (including pollen grains from flower tissues), and the supernatant was taken. This step was repeated until the supernatant cleared or a significant size of pellet was not observed. For root-containing tissues (seedlings), the nuclei suspension was filtered through Flowmi strainers (40 µm) to remove additional sources of debris. Finally, nuclei were pelleted by centrifugation at 700 x g for 10 min with a swing-rotor centrifuge and resuspended in an appropriate volume of NPB. Nuclei were counted manually with a hemocytometer. The nuclei suspension was loaded to 10x Genomics Chip M, which was then processed with 10x Genomics Chromium X controller, and sequencing libraries were constructed following manufacturer’s instructions. Each sequencing library was sequenced with Illumina NovaSeq 6000.

### Sample multiplexing

For the stem sample, three regions of the stem (base, branched area, and apex) were harvested separately and multiplexed using the 3’ CellPlex Kit following the manufacturer’s specifications (10x Genomics). Briefly, equal quantities of purified nuclei from each sample were individually labeled with separate CellPlex barcodes. Following labeling, the nuclei were pelleted and washed twice with PBS containing 2% BSA with RNase inhibitor. The nuclei were then quantified again by hemocytometer and were pooled in equal quantities and brought to the appropriate volume for loading into the 10x Genomics Chip M as described previously.

### snRNA-seq pre-processing

Demultiplexed FASTQ files were used to generate gene-by-cell matrices with Cell Ranger (v7.0.0) and were aligned to the Arabidopsis nuclear transcriptome using the – include-introns flag, which was prepared with the TAIR10 genome and Araport 11 transcriptome. Downstream analyses were performed with Seurat v4.1.0. Low-quality and potential doublet nuclei were filtered separately for the relaxed and stringent datasets. For relaxed filtering, nuclei with fewer than 300 detected genes, and/or greater than 5% mitochondrial or 15% chloroplastic reads were removed. For the stringently filtered dataset, nuclei that did not meet a cutoff of greater than 300 and fewer than 7,000 genes detected, > 400 UMI per nucleus, < 5% mitochondrial or <15% chloroplastic reads were removed.

### snRNA-seq analysis

For the analysis of single tissue datasets, after filtering 2,000 integration features for the aggregated datasets was determined for data scaling and dimensionality reduction using the SelectIntegrationFeatures function. Each of the datasets were then integrated using the IntegrateData function with 20 principal components (PCs). Following integration of each dataset, the entire dataset was scaled and dimensionality reduction was performed with PCA followed by UMAP using 30 PCs. Identification of nearest neighbors and clustering was then performed using the FindNeighbors and FindClusters functions, respectively. Cluster markers were then identified for each cluster using the FindAllMarkers function with logfc.theshold = 0.25 and min.pct = 0.1. For iterative subclustering, each individual cluster was subset and re-normalized, 2,000 variable features were identified, and the data was scaled, dimensions were reduced by PCA and UMAP using 30 PCs, and neighbors and clusters were identified.

For the globally integrated dataset, each of the individual stringently filtered datasets were merged, and the data was normalized. Following the guidelines of^1,2^, 4,000 variable features were identified and used for PCA, and pre clustering was performed with Harmony^3^ using 50 PCs. t-distributed stochastic neighbor embedding (t-SNE) was then performed with^4^ using the harmony embeddings with the perplexity value equal to the number of nuclei in the dataset / 100 and learning rate equal to the number of nuclei / 12.

### Cell type annotation

For cell annotation we used marker genes identified in previous studies using promoters GUS/GFP fusions, in situ hybridization assays, bulk RNA-seq studies of dissected tissues, single-cell atlases of roots, shoot apical meristem and inflorescence stem (Table S1). For *de novo*-identified marker genes of each cluster, the number of curated cell type markers was divided by the number of cluster marker genes to calculate the cell type enrichment score. We also explored the expression of top cluster markers in databases such as TAIR (www.arabidopsis.org) and ePlant (bar.utoronto.ca/eplant/).

Silique embryo:

For annotating silique major clusters, a curated list of marker genes were used^27^.

### Single-nucleus and single-cell dataset integration and cell type prediction

To integrate our 21d– and 30d-old rosette single-nuclei datasets with a published leaf protoplast dataset^24^, each dataset was individually loaded, normalized and scaled using SCTransform^32^, and 30 PCs were used for dimensionality reduction. Integration anchors were calculated using the FindTransferAnchors function using the protoplast dataset as the reference dataset. Cell type label transfer, low-dimensional embedding correction, and dataset projection of the nuclei-datasets was performed using the MapQuery function.

### Seedling and rosette dataset integration

To integrate the seedling and rosette time-series datasets, each dataset was pre-processed as described previously, and each dataset within the developmental time series was merged. For each merged dataset, Principal components were calculated, and Harmony^3^ was used to integrate the datasets regressing variables of dataset input, and % mitochondrial and chloroplastic reads, rounded to the nearest integer. Dimensionality reduction, nearest neighbor identification, and clustering was then performed using 20 harmony dimensions. Identification of cluster markers was performed as described previously for individual datasets.

### Identification of unique cell population marker genes

First, we identified marker genes for individual clusters of each tissue type (i.e., for 183 clusters) using the *FindAllMarkers* function of Seurat (adjusted p-value < 0.0005, average log_2_ fold change > 2). Then, we checked if these cluster marker genes are highly expressed in other organs using pseudobulk expression data of each cluster. A marker gene candidate was removed when (1) its normalized expression level in the target cluster was not more than four-fold higher than any other clusters in the other organs or (2) the log_2_ transcript per million (TPM) value of the gene is higher than five in any clusters in the other organs. In this way, we removed genes that are cell type-specific in a particular organ but are also expressed in other organs. In this analysis, we defined organs in the following way: imbibed and germinating seeds (0d and 1.25d), seedlings (3d, 6d, and 12d), rosettes (21d and 30d), stems, flowers, and siliques.

### Pseudotime analysis

To determine genes involved in root hair development, the annotated root hair clusters were subset and re-clustered following normalization, scaling and integration of the subsetted data. Cluster centroids were identified, trajectory inference and pseudotime estimation were performed using the Slingshot package^5^. Identification of genes with differential expression patterns along pseudotime was performed using the tradeSeq package^6^. Genes with the most significant Wald values were identified and subsetted based on hierarchical clustering for visualization. The expression profiles of groups of hierarchically clustered genes associated with early, middle, and late pseudotime intervals were subset and the binned aggregate expression values were displayed.

### Cross-tissue cell type analysis

Clusters identified from the global dataset that were well represented by several datasets were selected and individually subsetted. Re-clustering was performed as described previously. For cross-tissue comparisons of datasets, the tissue dataset that comprised the majority population of each subcluster was specified as the organ identity for each subcluster. Systematic identification of subcluster markers was performed, and unique and overlapping sets of subcluster marker genes were visualized using ComplexUpset^7^. Intersections of subcluster groups with fewer than 10 shared subcluster markers were filtered for visualization.

### TF analysis

A list of Arabidopsis TFs was downloaded from PlantTFDB (http://planttfdb.gao-lab.org/) to subset our single-cell data. Phylogenetic analysis was performed on Clustral Omega based on amino acid sequences of TFs. Phylogenetic trees were generated on Interactive Tree Of Life (iTOL).

### Tissue rank analysis (related to Fig 6B)

The major clusters in all tissues were ordered and ranked by expression of each TF (1 for the cluster with the lowest gene expression). Tissue ranks were calculated by taking the mean rank of clusters in each tissue for each TF.

### MERFISH panel design

A list of marker genes of silique major and sub clusters were filtered by the following criteria: (1) more than 25 specific probes can be designed based on Vizgen’s software and (2) the transcript per million (TPM) value is below 1000 in silique pseudobulk RNA-seq data (obtained by aggregating all the silique cells). After the filtering, 140 genes representing most major clusters and selected sub clusters were chosen from the remaining marker genes. The panel design was balanced so that the total TPM value of 140 transcripts was below 11,000 to avoid overcrowding of signal.

### Tissue cryosectioning, fixation, and mounting

Plants were grown according to the previously described methods. Fully elongated siliques matching the aforementioned developmental stage were excised and were immediately incubated in OCT (Fisher) for 5 min on ice. Following incubation, the siliques were placed into cryomolds (Sakura) which were filled with OCT and immediately frozen in an isopentane bath cooled by liquid N_2_. Tissue blocks were acclimated to –18°C in a pre-cooled cryostat chamber (Leica) for 1 h. Tissue blocks were trimmed until the tissue was entered, after which 10 µm sections were visually inspected until the region of interest was exposed. Sample mounting and preparation were performed according to the MERSCOPE user guide, with slight modifications. Briefly, a 10 µm section was melted and mounted onto a room temperature MERSCOPE slide (Vizgen, 20400001), placed into a 60 mm Petri dish and re-frozen by incubation in the cryostat chamber for 5 min. Subsequent steps were performed with the mounted samples in the Petri dish. The samples were then baked at 37°C for 5 min and were then incubated in fixation buffer (1X PBS, 4% formaldehyde) for 15 min at RT. Samples were then washed with 1X PBS containing 1:500 RNAse inhibitor (Protector RNAse inhibitor, Millipore Sigma) for 5 min at RT in triplicate. Following the final PBS wash, samples were dehydrated by incubation in 70% EtOH at 4°C overnight.

### MERFISH experiment

Tissue sections were processed following Vizgen’s protocol. After removing 70% ethanol, the sample was incubated in the Sample Prep Wash Buffer (PN20300001) for 1 min then incubated in the Formamide Wash Buffer (PN20300002) at 37°C for 30 min. After removing the Formamide Wash Buffer, the sample was incubated in MERSCOPE Gene Panel Mix at 37°C for 42 h. After the probe hybridization, the sample was washed twice with the Formamide Wash Buffer at 47°C for 30 min and once with the Sample Prep Wash Buffer at RT for 2 min. After the washing, the sample was embedded in hydrogel by incubating in the Gel Embedding Solution (Gel Embedding Premix (PN20300004), 10% w/v ammonium persulfate solution, and N,N,N’,N’-tetramethylethylenediamine) at RT for 1.5 h. Then, the sample was cleared by first incubating in the Digestion Mix (Digestion Premix (PN 20300005) and 1:40 RNase inhibitor) at RT for 2 h, followed by the incubation in the Clearing Solution (Clearing Premix (PN 20300003) and Proteinase K) at 47°C for 24 h then at 37°C for 24 h. The cleared sample was washed twice with the Sample Prep Wash Buffer and stained with DAPI and PolyT Staining Reagent at RT for 15 min then washed with the Formamide Wash Buffer at RT for 10 min and rinsed with the Sample Prep Wash Buffer. The sample was imaged with the MERSCOPE Instrument, and detected transcripts were decoded on the MERSCOPE Instrument using a Codebook generated by Vizgen. Transcripts were visualized on Vizgen’s Vizualizer.

### Curio Seeker spatial transcriptomics

Germinating seeds (1.25d) were grown as previously described and were subsequently submerged in OCT in a cryomold (Sakura), and flash frozen in a liquid N_2_ cooled isopentane bath. Cryosectioning of the 1.25d germinating seeds was performed as described above. Following sectioning, the frozen section was arranged onto the center of the Curio Seeker Spatial Mapping Kit (Curio Bioscience) and gently melted and stored at –80°C until further processing according to the manufacturer’s specifications. Briefly, the frozen tile was gently thawed to RT and submerged in Hybridization Reaction Mix for 15 min, followed by sequential transfers to wash buffer and RT Reaction Mix at RT. Reverse transcription was performed by incubation at 52°C for 30 min followed by tissue digestion with the addition of Tissue Clearing Reaction Mix and incubation at 37°C for 30 min. Following tissue digestion, the beads were dissociated from the tile by the addition of Bead Wash Buffer and pipetting. The released beads were transferred to a new tube and pelleted by centrifugation and washed twice in Bead Wash Buffer. Following the second wash, the beads were incubated at 95°C for 5 min and then pelleted and immediately resuspended Second Strand Mix and incubated at 37°C for 60 min. Following incubation, Bead Wash Buffer was added, and the beads were washed three times with Bead Wash Buffer. After the third wash, the beads were resuspended in and amplified in cDNA Amplification Mix by PCR. The amplified cDNA was then purified and quantified, followed by tagmentation and cleanup. The libraries were then sequenced on the NovaSeq 6000.

### Spatial transcriptomics analysis

Spot demultiplexing was performed according to the manufacturer’s specifications using the Seeker Bioinformatic Pipeline (Curio Bioscience). The resulting spot-by-gene matrix was utilized for downstream analysis with Seurat v4.1.0. Empty and low-quality spots with fewer than 125 genes detected or 125 UMIs were removed. The filtered dataset was normalized with SCTransform with the variable features calculated with a residual variance cutoff of 1. Principal components were calculated and used for UMAP dimensionality reduction with 20 PCs, followed by nearest neighbor identification and clustering. Identification of cluster markers was performed as described previously.

## Data availability

Raw sequencing data and processed data have been deposited at GEO: GSE226097

**Figure S1.**
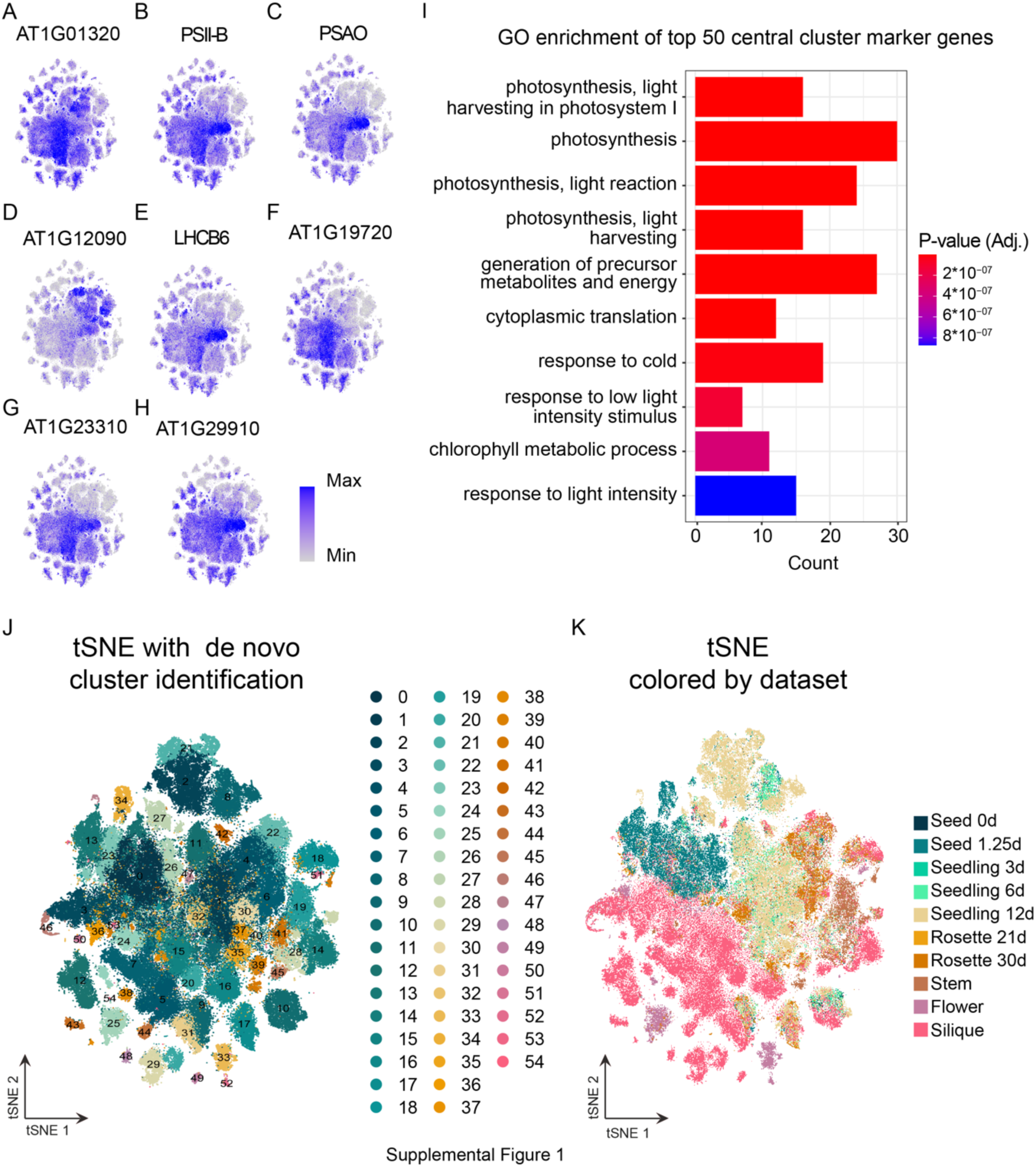
Characteristics of the central cluster. (**A-H**) Expression of representative genes with enriched expression in the central cluster in the globally integrated dataset. (**I**) GO term enrichment of the top 50 marker genes identified in the central cluster. (**J and K**) Dimensionality reduction and clustering of the globally integrated dataset with the large central cluster removed, colored by (J) *de novo* cluster identification and (K) colored by dataset.

**Figure S2.**
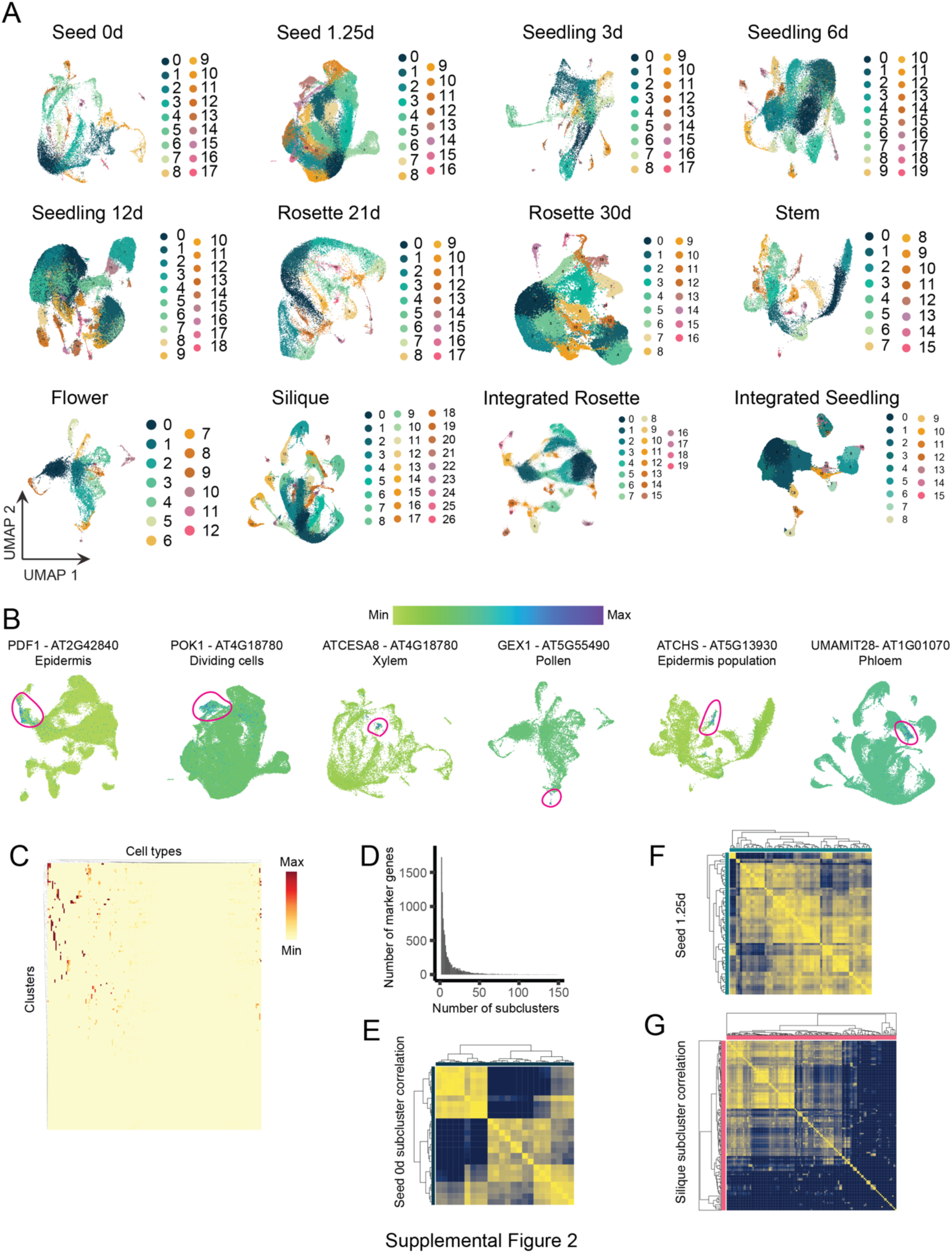
Clustering of individual datasets. (**A**) Dimensionality reduction and *de novo* clustering results for each individual dataset. (**B**) Expression patterns of known cell type-specific marker genes as visualized in our Arabidopsis developmental atlas browser. Regions of enriched expression are circled. (**C**) Marker gene enrichment score for each cell type/state in each major cluster. (**D**) Histogram showing the number of *de novo* identified subclusters marker genes that are shared between subclusters. The x axis shows the number of subclusters that share the same marker gene. (**E-G**) Correlation of pseudobulk expression between subclusters of the (E) 0d seed, (F) 1.25d seed, and (F) silique tissues.

**Figure S3.**
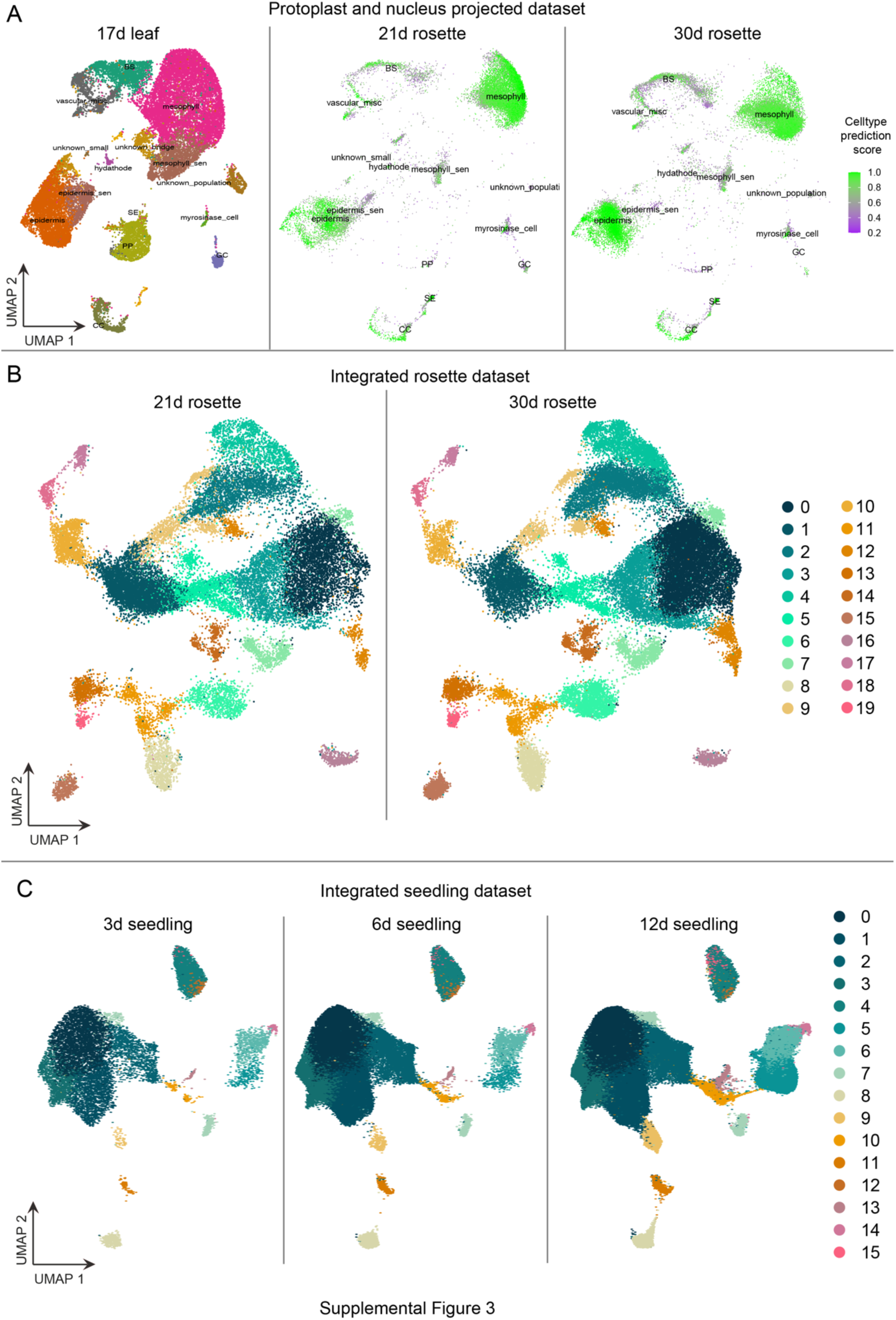
Projection and integration of time series datasets. (**A**) Cell type prediction scores of the 21d-(middle) and 30d-old (right) rosette nuclei datasets when projected onto a leaf protoplast dataset (left). (**B**) Integration of the 21d-(left) and 30d-old (right) rosette nuclei datasets. (**C**) Integration of the 3d-, 6d-, and 12d-old seedling datasets.

**Figure S4.**
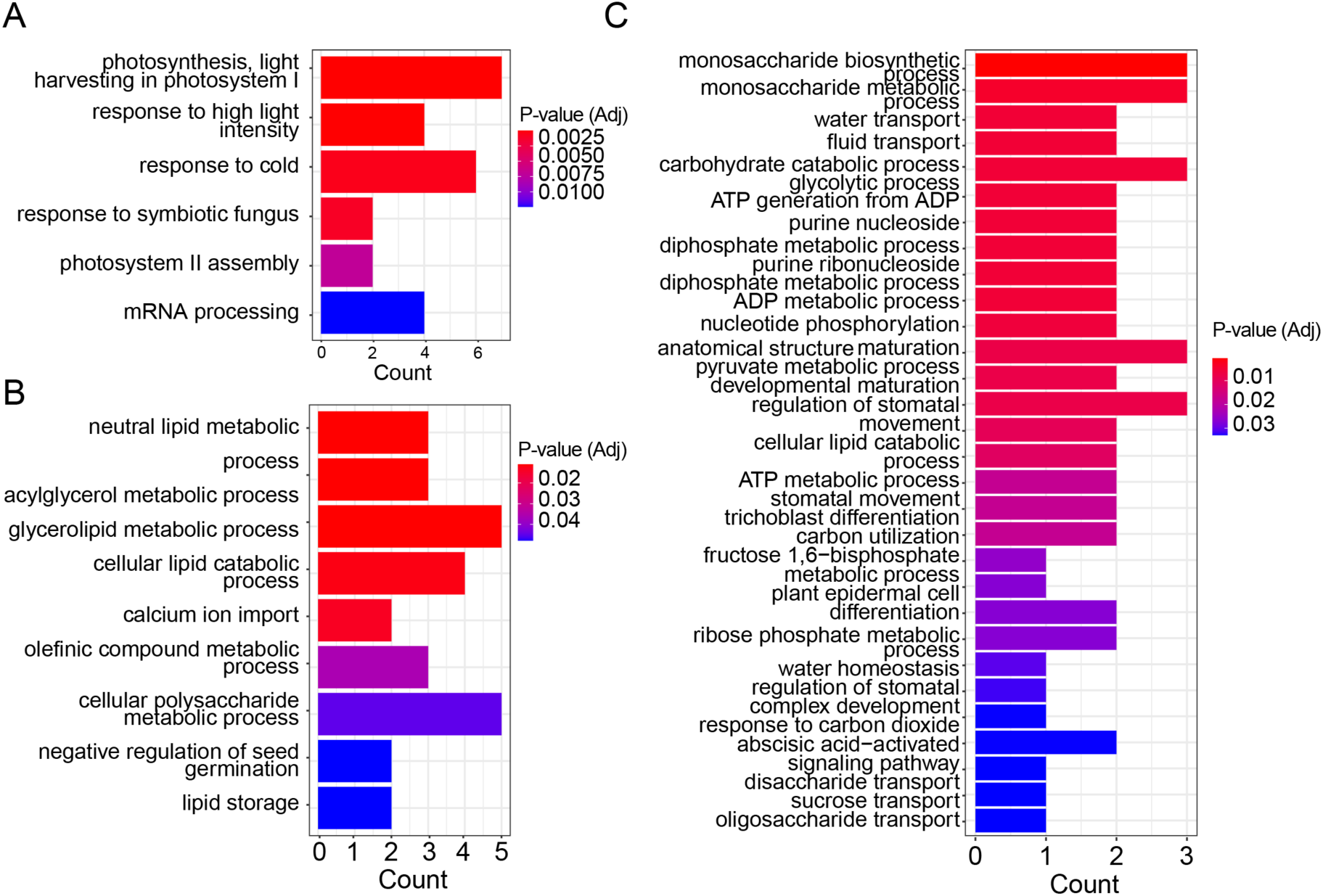
GO terms of root hair pseudotime expression modules. (**A** to **C**) GO term enrichment of genes differentially regulated over pseudotime of root hair cells corresponding to (A) root hair initiation, (B) root hair differentiation, and (C) root hair maturation.

**Figure S5.**
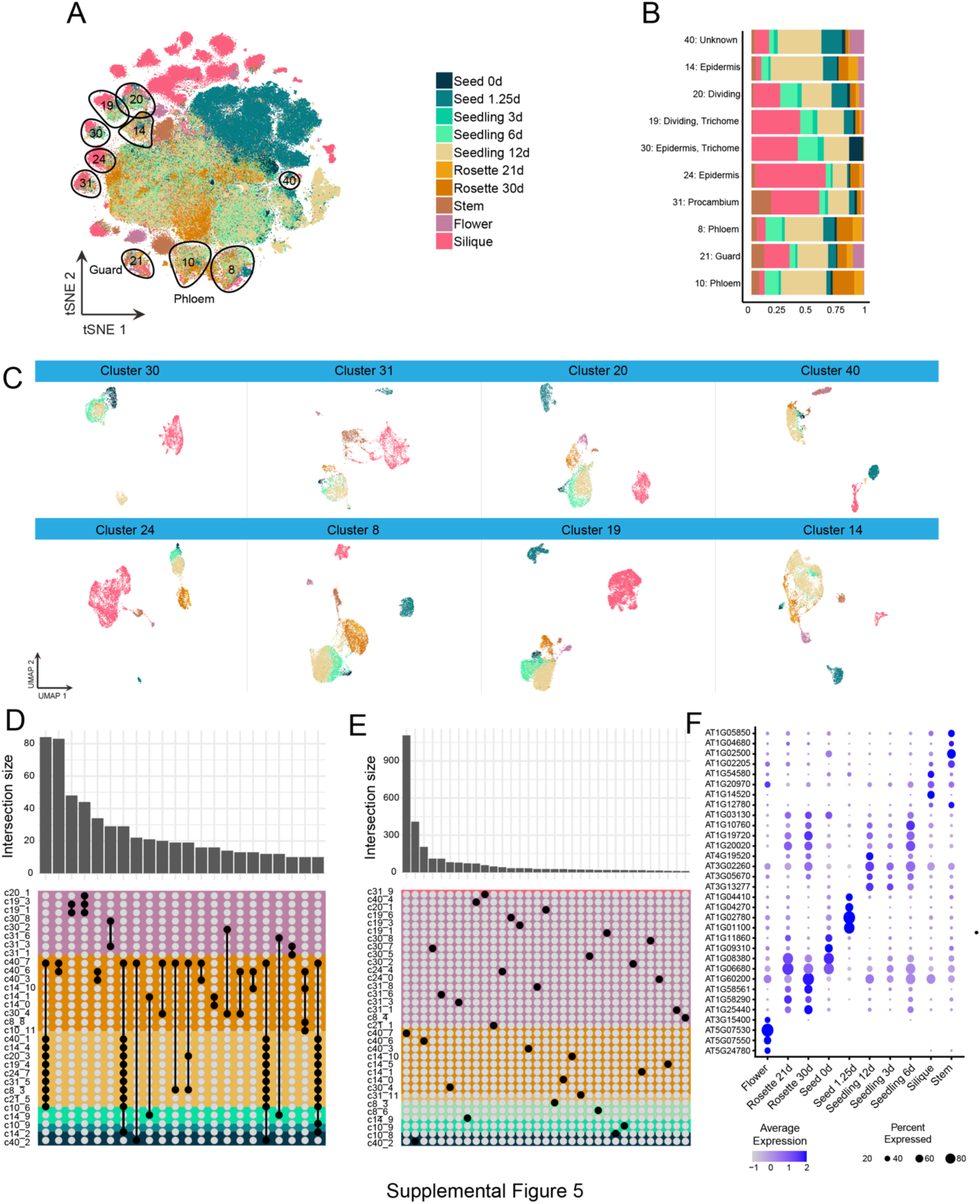
Cross-tissue cell type analysis of ten cell type populations. (**A**) The ten cell populations included in the cross-cell type analysis highlighted in the globally clustered tSNE. (**B**) Proportion of cells per dataset for the ten clusters analyzed. (**C**) Subclustering of the remaining eight clusters included in the ten-cluster analysis. Nuclei are colored by dataset of origin. (**D**) Quantification of identified markers shared between subcluster populations. Groups are named by cluster of origin and subcluster number. Tissue of origin is depicted by the colored bar. Subcluster intersections with greater than ten shared markers are depicted. (**E**) Quantification of subcluster markers uniquely identified as a marker in only one subcluster population. Groups are named by cluster of origin and subcluster number. Tissue of origin is depicted by the colored bar. Subclusters with greater than ten uniquely identified marker genes are depicted. (**F**) Expression of representative tissue-level markers identified for each tissue.

**Figure S6.**
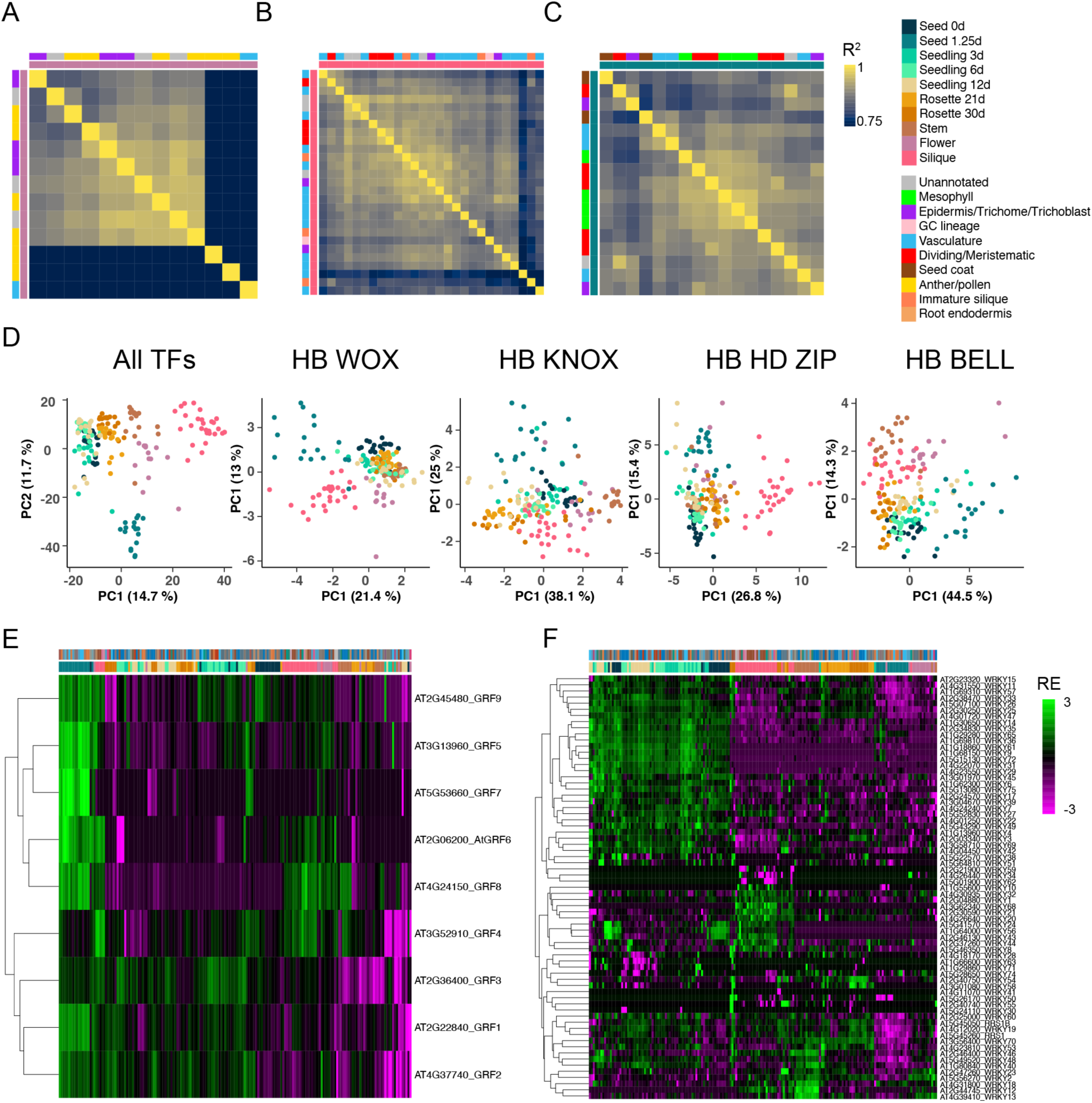
Transcription factor (TF) analysis. (**A-C**) Heatmaps showing pair-wise correlation of pseudobulk TF expression between major clusters of flower (left), silique (middle), and seed 1.25d (right). (**D**) Principal component analysis of TF family expression in each cluster. Each dot represents a cluster colored by the tissue of origin. (**E** and **F**) Heatmaps showing expression of GRF (left) and WRKY (right) family TFs in all major clusters. Expression was shown as log2 relative expression (RE).

**Figure S7.**
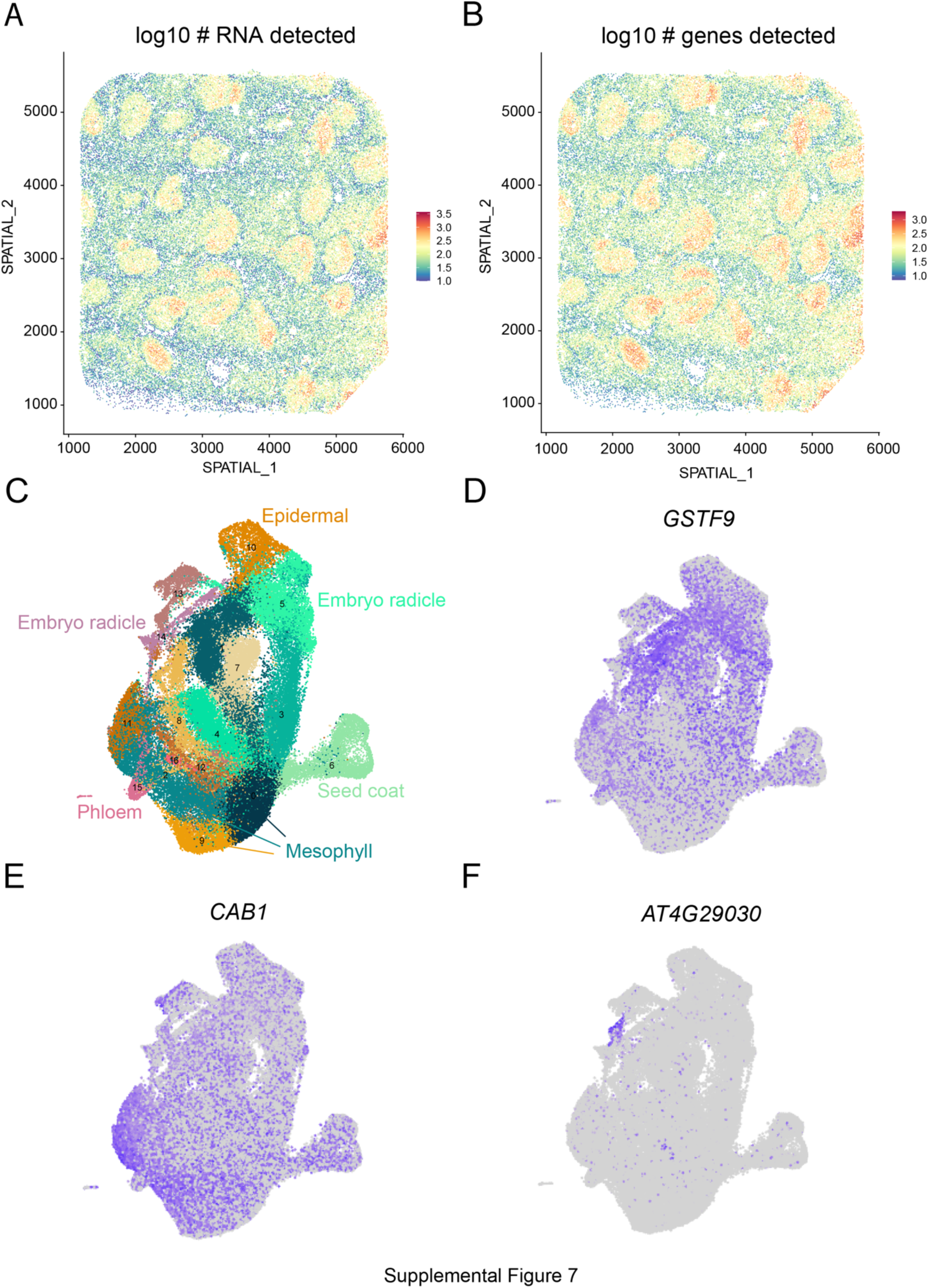
Spatial transcriptomics of the 1.25d germinating seed. (**A** and **B**) Quality metrics of the spatial transcriptomics dataset. Log_10_ values of (A) the number of transcripts and (B) the number of genes detected per spot in spatial coordinates. (**C-F**) Cross-comparison of spatial transcriptomics derived cluster markers in the matched droplet-based single nucleus dataset. (C) Dimensionality reduction and clustering of the droplet-based 1.25d seed dataset in Figure 2A. (D-F) Expression of cluster markers identified in the spatial transcriptomics dataset corresponding to the (D) stele (cluster 4), (E) cotyledon (cluster 5), and (F) unannotated (cluster 6) in the droplet-based dataset.

## Table captions

Table S1. Cell type and tissue marker genes collected from the literature and this study

Table S2. Table of cluster markers and cell type annotation of individual tissue datasets

Table S3. Pseudobulk expression values of 183 major clusters

Table S4. Pseudobulk expression values of 655 subclusters

Table S5. Genes differentially expressed over pseudotime of root hair development

Table S7. List of transcription factors identified as cluster markers from the 183 major clusters

## References

1. Larkin, J.C., Brown, M.L., and Schiefelbein, J. (2003). How do cells know what they want to be when they grow up? Lessons from epidermal patterning in Arabidopsis. Annu. Rev. Plant Biol. 54, 403–430.

2. De Rybel, B., Mähönen, A.P., Helariutta, Y., and Weijers, D. (2016). Plant vascular development: from early specification to differentiation. Nat. Rev. Mol. Cell Biol. 17, 30–40.

3. Meyer, H.M., Teles, J., Formosa-Jordan, P., Refahi, Y., San-Bento, R., Ingram, G., Jönsson, H., Locke, J.C.W., and Roeder, A.H.K. (2017). Fluctuations of the transcription factor ATML1 generate the pattern of giant cells in the Arabidopsis sepal. Elife 6. 10.7554/eLife.19131.

4. Roeder, A.H.K., Chickarmane, V., Cunha, A., Obara, B., Manjunath, B.S., and Meyerowitz, E.M. (2010). Variability in the control of cell division underlies sepal epidermal patterning in Arabidopsis thaliana. PLoS Biol. 8, e1000367.

5. Ripoll, J.-J., Zhu, M., Brocke, S., Hon, C.T., Yanofsky, M.F., Boudaoud, A., and Roeder, A.H.K. (2019). Growth dynamics of the Arabidopsis fruit is mediated by cell expansion. Proc. Natl. Acad. Sci. U. S. A. 116, 25333–25342.

6. Domcke, S., and Shendure, J. (2023). A reference cell tree will serve science better than a reference cell atlas. Cell 186, 1103–1114.

7. Menges, M., Hennig, L., Gruissem, W., and Murray, J.A.H. (2002). Cell cycle-regulated gene expression inArabidopsis. J. Biol. Chem. 277, 41987–42002.

8. Lu, T.-C., Brbić, M., Park, Y.-J., Jackson, T., Chen, J., Kolluru, S.S., Qi, Y., Katheder, N.S., Cai, X.T., Lee, S., et al. (2023). Aging Fly Cell Atlas identifies exhaustive aging features at cellular resolution. Science 380, eadg0934.

9. Qi, F., and Zhang, F. (2019). Cell Cycle Regulation in the Plant Response to Stress. Front. Plant Sci. 10, 1765.

10. Cole, B., Bergmann, D., Blaby-Haas, C.E., Blaby, I.K., Bouchard, K.E., Brady, S.M., Ciobanu, D., Coleman-Derr, D., Leiboff, S., Mortimer, J.C., et al. (2021). Plant single-cell solutions for energy and the environment. Commun Biol 4, 962.

11. Taylor, I.W., Rahul Patharkar, O., Hsu, C.-W., Baer, J., Niederhuth, C.E., Ohler, U., Benfey, P.N., and Walker, J.C. (2022). Single-cell transcriptomics of the Arabidopsis floral abscission zone. bioRxiv. 10.1101/2022.07.14.500021.

12. Zhang, T.-Q., Chen, Y., and Wang, J.-W. (2021). A single-cell analysis of the Arabidopsis vegetative shoot apex. Dev. Cell 56, 1056–1074.e8.

13. Song, Q., Ando, A., Jiang, N., Ikeda, Y., and Chen, Z.J. (2020). Single-cell RNA-seq analysis reveals ploidy-dependent and cell-specific transcriptome changes in Arabidopsis female gametophytes. Genome Biol. 21, 178.

14. Apelt, F., Mavrothalassiti, E., Gupta, S., Machin, F., Olas, J.J., Annunziata, M.G., Schindelasch, D., and Kragler, F. (2022). Shoot and root single cell sequencing reveals tissue– and daytime-specific transcriptome profiles. Plant Physiol. 188, 861–878.

15. Misra, C.S., Santos, M.R., Rafael-Fernandes, M., Martins, N.P., Monteiro, M., and Becker, J.D. (2019). Transcriptomics of Arabidopsis sperm cells at single-cell resolution. Plant Reprod. 32, 29–38.

16. Shahan, R., Hsu, C.-W., Nolan, T.M., Cole, B.J., Taylor, I.W., Greenstreet, L., Zhang, S., Afanassiev, A., Vlot, A.H.C., Schiebinger, G., et al. (2022). A single-cell Arabidopsis root atlas reveals developmental trajectories in wild-type and cell identity mutants. Developmental Cell 57, 543–560. 10.1016/j.devcel.2022.01.008.

17. Shulse, C.N., Cole, B.J., Ciobanu, D., Lin, J., Yoshinaga, Y., Gouran, M., Turco, G.M., Zhu, Y., O’Malley, R.C., Brady, S.M., et al. (2019). High-Throughput Single-Cell Transcriptome Profiling of Plant Cell Types. Cell Rep. 27, 2241–2247.e4.

18. Ryu, K.H., Huang, L., Kang, H.M., and Schiefelbein, J. (2019). Single-Cell RNA Sequencing Resolves Molecular Relationships Among Individual Plant Cells. Plant Physiol. 179, 1444–1456.

19. Nobori, T., Monell, A., Lee, T.A., Zhou, J., Nery, J., and Ecker, J.R. (2023). Time-resolved single-cell and spatial gene regulatory atlas of plants under pathogen attack. bioRxiv, 2023.04.10.536170. 10.1101/2023.04.10.536170.

20. Boyes, D.C., Zayed, A.M., Ascenzi, R., McCaskill, A.J., Hoffman, N.E., Davis, K.R., and Görlach, J. (2001). Growth Stage–Based Phenotypic Analysis of Arabidopsis: A Model for High Throughput Functional Genomics in Plants. Plant Cell 13, 1499– 1510.

21. Mizzotti, C., Rotasperti, L., Moretto, M., Tadini, L., Resentini, F., Galliani, B.M., Galbiati, M., Engelen, K., Pesaresi, P., and Masiero, S. (2018). Time-Course Transcriptome Analysis of Arabidopsis Siliques Discloses Genes Essential for Fruit Development and Maturation. Plant Physiol. 178, 1249–1268.

22. Alvarez-Buylla, E.R., Benítez, M., Corvera-Poiré, A., Chaos Cador, A., de Folter, S., Gamboa de Buen, A., Garay-Arroyo, A., García-Ponce, B., Jaimes-Miranda, F., Pérez-Ruiz, R.V., et al. (2010). Flower development. Arabidopsis Book 8, e0127.

23. Li, H., Janssens, J., De Waegeneer, M., Kolluru, S.S., Davie, K., Gardeux, V., Saelens, W., David, F., Brbić, M., Leskovec, J., et al. Fly Cell Atlas: a single-cell transcriptomic atlas of the adult fruit fly. Science 375. 10.1101/2021.07.04.451050.

24. Lopez-Anido, C.B., Vatén, A., Smoot, N.K., Sharma, N., Guo, V., Gong, Y., Anleu Gil, M.X., Weimer, A.K., and Bergmann, D.C. (2021). Single-cell resolution of lineage trajectories in the Arabidopsis stomatal lineage and developing leaf. Dev. Cell 56, 1043–1055.e4.

25. Procko, C., Lee, T., Borsuk, A., Bargmann, B.O.R., Dabi, T., Nery, J.R., Estelle, M., Baird, L., O’Connor, C., Brodersen, C., et al. (2022). Leaf cell-specific and single-cell transcriptional profiling reveals a role for the palisade layer in UV light protection. Plant Cell 34, 3261–3279.

26. Kim, J.-Y., Symeonidi, E., Pang, T.Y., Denyer, T., Weidauer, D., Bezrutczyk, M., Miras, M., Zöllner, N., Hartwig, T., Wudick, M.M., et al. (2021). Distinct identities of leaf phloem cells revealed by single cell transcriptomics. Plant Cell 33, 511–530.

27. Denyer, T., Ma, X., Klesen, S., Scacchi, E., Nieselt, K., and Timmermans, M.C.P. (2019). Spatiotemporal Developmental Trajectories in the Arabidopsis Root Revealed Using High-Throughput Single-Cell RNA Sequencing. Dev. Cell 48, 840– 852.e5.

28. Picard, C.L., Povilus, R.A., Williams, B.P., and Gehring, M. (2021). Transcriptional and imprinting complexity in Arabidopsis seeds at single-nucleus resolution. Nat Plants 7, 730–738.

29. Berardini, T.Z., Reiser, L., Li, D., Mezheritsky, Y., Muller, R., Strait, E., and Huala, E. (2015). The Arabidopsis information resource: Making and mining the “gold standard” annotated reference plant genome. Genesis 53, 474–485.

30. Fucile, G., Di Biase, D., Nahal, H., La, G., Khodabandeh, S., Chen, Y., Easley, K., Christendat, D., Kelley, L., and Provart, N.J. (2011). ePlant and the 3D data display initiative: integrative systems biology on the world wide web. PLoS One 6, e15237.

31. Waese, J., Fan, J., Pasha, A., Yu, H., Fucile, G., Shi, R., Cumming, M., Kelley, L.A., Sternberg, M.J., Krishnakumar, V., et al. (2017). ePlant: Visualizing and Exploring Multiple Levels of Data for Hypothesis Generation in Plant Biology. Plant Cell 29, 1806–1821.

32. Gómez-Felipe, A., Marconi, M., Branchini, E., Wang, B., Bertrand-Rakusova, H., Stan, T., Burkiewicz, J., de Folter, S., Routier-Kierzkowska, A.-L., Wabnik, K., et al. (2023). Competing differentiation gradients coordinate fruit morphogenesis. bioRxiv. 10.1101/2023.01.19.524793.

33. Stuart, T., Butler, A., Hoffman, P., Hafemeister, C., Papalexi, E., Mauck, W.M., 3rd, Hao, Y., Stoeckius, M., Smibert, P., and Satija, R. (2019). Comprehensive Integration of Single-Cell Data. Cell 177, 1888–1902.e21.

34. Guillotin, B., Rahni, R., Passalacqua, M., Mohammed, M.A., Xu, X., Raju, S.K., Ramírez, C.O., Jackson, D., Groen, S.C., Gillis, J., et al. (2023). A pan-grass transcriptome reveals patterns of cellular divergence in crops. Nature 617, 785– 791.

35. Kim, J., Kim, J.H., Lyu, J.I., Woo, H.R., and Lim, P.O. (2018). New insights into the regulation of leaf senescence in Arabidopsis. J. Exp. Bot. 69, 787–799.

36. Woo, H.R., Koo, H.J., Kim, J., Jeong, H., Yang, J.O., Lee, I.H., Jun, J.H., Choi, S.H., Park, S.J., Kang, B., et al. (2016). Programming of Plant Leaf Senescence with Temporal and Inter-Organellar Coordination of Transcriptome in Arabidopsis. Plant Physiol. 171, 452–467.

37. Grierson, C., Nielsen, E., Ketelaarc, T., and Schiefelbein, J. (2014). Root hairs. Arabidopsis Book 12, e0172.

38. Stadler, R., and Sauer, N. (1996). TheArabidopsis thaliana AtSUC2Gene is Specifically Expressed in Companion Cells. Bot. Acta 109, 299–306.

39. Truernit, E., and Sauer, N. (1995). The promoter of the Arabidopsis thaliana SUC2 sucrose-H+ symporter gene directs expression of beta-glucuronidase to the phloem: evidence for phloem loading and unloading by SUC2. Planta 196, 564– 570.

40. Ohashi-Ito, K., and Bergmann, D.C. (2006). Arabidopsis FAMA controls the final proliferation/differentiation switch during stomatal development. Plant Cell 18, 2493–2505.

41. Bai, B., Peviani, A., van der Horst, S., Gamm, M., Snel, B., Bentsink, L., and Hanson, J. (2017). Extensive translational regulation during seed germination revealed by polysomal profiling. New Phytol. 214, 233–244.

42. Liew, L.C., You, Y., Oliva, M., Peirats-Llobet, M., Ng, S., Tamiru-Oli, M., Berkowitz, O., Hong, U.V.T., Haslem, A., Stuart, T., et al. (2023). Establishment of cell transcriptional identity during seed germination. bioRxiv, 2023.01.21.523180. 10.1101/2023.01.21.523180.

43. Peirats-Llobet, M., Yi, C., Liew, L.C., Berkowitz, O., Narsai, R., Lewsey, M.G., and Whelan, J. (2023). Spatially resolved transcriptomic analysis of the germinating barley grain. Nucleic Acids Res. 51, 7798–7819.

44. Browning, K.S., and Bailey-Serres, J. (2015). Mechanism of cytoplasmic mRNA translation. Arabidopsis Book 13, e0176.

45. Jakoby, M., Weisshaar, B., Dröge-Laser, W., Vicente-Carbajosa, J., Tiedemann, J., Kroj, T., Parcy, F., and bZIP Research Group (2002). bZIP transcription factors in Arabidopsis. Trends Plant Sci. 7, 106–111.

46. Li, M., Lin, W., Hinckley, W., Yao, T., Muchero, W., Chen, J.-G., and Carol Huang, S. (2022). Uncovering the “ZIP code” for bZIP dimers reveals novel motifs, regulatory rules and one billion years of cis-element evolution. bioRxiv. 10.1101/2022.04.17.488518.

47. De Smet, R., Sabaghian, E., Li, Z., Saeys, Y., and Van de Peer, Y. (2017). Coordinated Functional Divergence of Genes after Genome Duplication in Arabidopsis thaliana. Plant Cell 29, 2786–2800.

48. Stickels, R.R., Murray, E., Kumar, P., Li, J., Marshall, J.L., Di Bella, D.J., Arlotta, P., Macosko, E.Z., and Chen, F. (2021). Highly sensitive spatial transcriptomics at near-cellular resolution with Slide-seqV2. Nat. Biotechnol. 39, 313–319.

49. Olas, J.J., Apelt, F., Watanabe, M., Hoefgen, R., and Wahl, V. (2021). Developmental stage-specific metabolite signatures in Arabidopsis thaliana under optimal and mild nitrogen limitation. Plant Sci. 303, 110746.

50. Tohge, T., Borghi, M., and Fernie, A.R. (2018). The natural variance of the Arabidopsis floral secondary metabolites. Sci Data 5, 180051.

51. Li, H., Janssens, J., Waegeneer, M.D., Kolluru, S.S., Davie, K., Gardeux, V., Saelens, W., David, F.P.A., Brbić, M., Spanier, K., et al. (2022). Fly Cell Atlas: A single-nucleus transcriptomic atlas of the adult fruit fly. Science 375, eabk2432.

52. Nobori, T., Oliva, M., Lister, R., and Ecker, J.R. (2023). Multiplexed single-cell 3D spatial gene expression analysis in plant tissue using PHYTOMap. Nature Plants, 1–8.

## Supplemental References

53. Liu, H., Zhou, J., Tian, W., Luo, C., Bartlett, A., Aldridge, A., Lucero, J., Osteen, J.K., Nery, J.R., Chen, H., et al. (2021). DNA methylation atlas of the mouse brain at single-cell resolution. Nature 598, 120–128.

54. Kobak, D., and Berens, P. (2019). The art of using t-SNE for single-cell transcriptomics. Nat. Commun. 10, 5416.

55. Korsunsky, I., Millard, N., Fan, J., Slowikowski, K., Zhang, F., Wei, K., Baglaenko, Y., Brenner, M., Loh, P.-R., and Raychaudhuri, S. (2019). Fast, sensitive and accurate integration of single-cell data with Harmony. Nat. Methods 16, 1289–1296.

56. Linderman, G.C., Rachh, M., Hoskins, J.G., Steinerberger, S., and Kluger, Y. (2019). Fast interpolation-based t-SNE for improved visualization of single-cell RNA-seq data. Nat. Methods 16, 243–245.

57. Street, K., Risso, D., Fletcher, R.B., Das, D., Ngai, J., Yosef, N., Purdom, E., and Dudoit, S. (2018). Slingshot: cell lineage and pseudotime inference for single-cell transcriptomics. BMC Genomics 19, 477.

58. Van den Berge, K., Roux de Bézieux, H., Street, K., Saelens, W., Cannoodt, R., Saeys, Y., Dudoit, S., and Clement, L. (2020). Trajectory-based differential expression analysis for single-cell sequencing data. Nat. Commun. 11, 1201.

59. Krassowski, M., Arts, M., Lagger, C., and Max (2022). krassowski/complex-upset: v1.3.5 (Zenodo) 10.5281/ZENODO.3700590.

60. Alandete-Saez, M., Ron, M., & McCormick, S. (2008). GEX3, expressed in the male gametophyte and in the egg cell of Arabidopsis thaliana, is essential for micropylar pollen tube guidance and plays a role during early embryogenesis. Molecular plant, 1(4), 586–598.

61. Barth, C., & Jander, G. (2006). Arabidopsis myrosinases TGG1 and TGG2 have redundant function in glucosinolate breakdown and insect defense. The Plant journal: for cell and molecular biology, 46(4), 549–562.

62. Borges, F., Gardner, R., Lopes, T., Calarco, J. P., Boavida, L. C., Slotkin, R. K., Martienssen, R. A., & Becker, J. D. (2012). FACS-based purification of Arabidopsis microspores, sperm cells and vegetative nuclei. Plant methods, 8(1), 44.

63. Endo, M., Mochizuki, N., Suzuki, T., & Nagatani, A. (2007). CRYPTOCHROME2 in vascular bundles regulates flowering in Arabidopsis. The Plant cell, 19(1), 84–93.

64. Endo, M., Shimizu, H., Nohales, M. A., Araki, T., & Kay, S. A. (2014). Tissue-specific clocks in Arabidopsis show asymmetric coupling. Nature, 515(7527), 419–422.

65. Engel, M. L., Holmes-Davis, R., & McCormick, S. (2005). Green sperm. Identification of male gamete promoters in Arabidopsis. Plant physiology, 138(4), 2124–2133.

66. Glaring, M. A., Zygadlo, A., Thorneycroft, D., Schulz, A., Smith, S. M., Blennow, A., & Baunsgaard, L. (2007). An extra-plastidial alpha-glucan, water dikinase from Arabidopsis phosphorylates amylopectin in vitro and is not necessary for transient starch degradation. Journal of experimental botany, 58(14), 3949–3960.

67. Guo, W. J., Bundithya, W., & Goldsbrough, P. B. (2003). Characterization of the Arabidopsis metallothionein gene family: tissue-specific expression and induction during senescence and in response to copper. The New phytologist, 159(2), 369– 381.

68. Kevei, Z., Baloban, M., Da Ines, O., Tiricz, H., Kroll, A., Regulski, K., Mergaert, P., & Kondorosi, E. (2011). Conserved CDC20 cell cycle functions are carried out by two of the five isoforms in Arabidopsis thaliana. PloS one, 6(6), e20618.

69. Kurata, T., Kawabata-Awai, C., Sakuradani, E., Shimizu, S., Okada, K., & Wada, T. (2003). The YORE-YORE gene regulates multiple aspects of epidermal cell differentiation in Arabidopsis. The Plant journal: for cell and molecular biology, 36(1), 55–66.

70. Long, J., Walker, J., She, W., Aldridge, B., Gao, H., Deans, S., Vickers, M., & Feng, X. (2021). Nurse cell--derived small RNAs define paternal epigenetic inheritance in Arabidopsis. Science (New York, N.Y.), 373(6550), eabh0556.

71. Lu, L., Lee, Y. R., Pan, R., Maloof, J. N., & Liu, B. (2005). An internal motor kinesin is associated with the Golgi apparatus and plays a role in trichome morphogenesis in Arabidopsis. Molecular biology of the cell, 16(2), 811–823.

72. Ma, X., Song, L., Yang, Y., & Liu, D. (2013). A gain-of-function mutation in the ROC1 gene alters plant architecture in Arabidopsis. The New phytologist, 197(3), 751–762.

73. Mathur, J., Molnár, G., Fujioka, S., Takatsuto, S., Sakurai, A., Yokota, T., Adam, G., Voigt, B., Nagy, F., Maas, C., Schell, J., Koncz, C., & Szekeres, M. (1998). Transcription of the Arabidopsis CPD gene, encoding a steroidogenic cytochrome P450, is negatively controlled by brassinosteroids. The Plant journal: for cell and molecular biology, 14(5), 593–602.

74. Mitchell, J., Mukhtar, N. K., Skinner, I., Bassel, G. W. (2018) Gibberellin response in the embryo epidermis regulates germination uniformity in response to seed priming. bioRxiv doi:10.1101/436121.

75. Müller, S., Han, S., & Smith, L. G. (2006). Two kinesins are involved in the spatial control of cytokinesis in Arabidopsis thaliana. Current biology: CB, 16(9), 888–894.

76. Nakayama, N., Arroyo, J. M., Simorowski, J., May, B., Martienssen, R., & Irish, V. F. (2005). Gene trap lines define domains of gene regulation in Arabidopsis petals and stamens. The Plant cell, 17(9), 2486–2506.

77. Okumoto, S., Schmidt, R., Tegeder, M., Fischer, W. N., Rentsch, D., Frommer, W. B., & Koch, W. (2002). High affinity amino acid transporters specifically expressed in xylem parenchyma and developing seeds of Arabidopsis. The Journal of biological chemistry, 277(47), 45338–45346.

78. Pilot, G., Stransky, H., Bushey, D. F., Pratelli, R., Ludewig, U., Wingate, V. P., & Frommer, W. B. (2004). Overexpression of GLUTAMINE DUMPER1 leads to hypersecretion of glutamine from Hydathodes of Arabidopsis leaves. The Plant cell, 16(7), 1827–1840.

79. Pommerrenig, B., Feussner, K., Zierer, W., Rabinovych, V., Klebl, F., Feussner, I., & Sauer, N. (2011). Phloem-specific expression of Yang cycle genes and identification of novel Yang cycle enzymes in Plantago and Arabidopsis. The Plant cell, 23(5), 1904–1919.

80. Qiu, J. L., Jilk, R., Marks, M. D., & Szymanski, D. B. (2002). The Arabidopsis SPIKE1 gene is required for normal cell shape control and tissue development. The Plant cell, 14(1), 101–118.

81. Ranjan, A., Fiene, G., Fackendahl, P., & Hoecker, U. (2011). The Arabidopsis repressor of light signaling SPA1 acts in the phloem to regulate seedling de-etiolation, leaf expansion and flowering time. Development (Cambridge, England), 138(9),

82. Rouse, D. T., Marotta, R., & Parish, R. W. (1996). Promoter and expression studies on an Arabidopsis thaliana dehydrin gene. FEBS letters, 381(3), 252–256.

83. Sawchuk, M. G., Donner, T. J., Head, P., & Scarpella, E. (2008). Unique and overlapping expression patterns among members of photosynthesis-associated nuclear gene families in Arabidopsis. Plant physiology, 148(4), 1908–1924.

84. Siegfried, K. R., Eshed, Y., Baum, S. F., Otsuga, D., Drews, G. N., & Bowman, J. L. (1999). Members of the YABBY gene family specify abaxial cell fate in Arabidopsis. Development (Cambridge, England), 126(18), 4117–4128.

85. Sistrunk, M. L., Antosiewicz, D. M., Purugganan, M. M., & Braam, J. (1994). Arabidopsis TCH3 encodes a novel Ca2+ binding protein and shows environmentally induced and tissue-specific regulation. The Plant cell, 6(11), 1553– 1565.

86. Takada, S., Takada, N., & Yoshida, A. (2013). ATML1 promotes epidermal cell differentiation in Arabidopsis shoots. Development (Cambridge, England), 140(9),

87. Uemoto, K., Araki, T., & Endo, M. (2018). Isolation of Arabidopsis Palisade and Spongy Mesophyll Cells. Methods in molecular biology (Clifton, N.J.), 1830, 141– 148.

88. Werner, T., Motyka, V., Laucou, V., Smets, R., Van Onckelen, H., & Schmülling, T. (2003). Cytokinin-deficient transgenic Arabidopsis plants show multiple developmental alterations indicating opposite functions of cytokinins in the regulation of shoot and root meristem activity. The Plant cell, 15(11), 2532–2550.

89. Wu, H., Li, L., Du, J., Yuan, Y., Cheng, X., & Ling, H. Q. (2005). Molecular and biochemical characterization of the Fe(III) chelate reductase gene family in Arabidopsis thaliana. Plant & cell physiology, 46(9), 1505–1514.

90. Yephremov, A., Wisman, E., Huijser, P., Huijser, C., Wellesen, K., & Saedler, H. (1999). Characterization of the FIDDLEHEAD gene of Arabidopsis reveals a link between adhesion response and cell differentiation in the epidermis. The Plant cell, 11(11), 2187–2201.

91. Zhang, X., Dyachok, J., Krishnakumar, S., Smith, L. G., & Oppenheimer, D. G. (2005). IRREGULAR TRICHOME BRANCH1 in Arabidopsis encodes a plant homolog of the actin-related protein2/3 complex activator Scar/WAVE that regulates actin and microtubule organization. The Plant cell, 17(8), 2314–2326.

92. Zhu, Y., Liu, L., Shen, L., & Yu, H. (2016). NaKR1 regulates long-distance movement of FLOWERING LOCUS T in Arabidopsis. Nature plants, 2(6), 16075.

93. Zimmermann, I., Saedler, R., Mutondo, M., & Hulskamp, M. (2004). The Arabidopsis GNARLED gene encodes the NAP125 homolog and controls several actin-based cell shape changes. Molecular genetics and genomics: MGG, 272(3), 290–296.

94. Abe, M., Takahashi, T., & Komeda, Y. (1999). Cloning and characterization of an L1 layer-specific gene in Arabidopsis thaliana. Plant & cell physiology, 40(6), 571–580.

95. Abe, M., Takahashi, T., & Komeda, Y. (2001). Identification of a cis-regulatory element for L1 layer-specific gene expression, which is targeted by an L1-specific homeodomain protein. The Plant journal: for cell and molecular biology, 26(5), 487–

96. Baima, S., Nobili, F., Sessa, G., Lucchetti, S., Ruberti, I., & Morelli, G. (1995). The expression of the Athb-8 homeobox gene is restricted to provascular cells in Arabidopsis thaliana. Development (Cambridge, England), 121(12), 4171–4182.

97. Banerjee, J., Sahoo, D. K., Dey, N., Houtz, R. L., & Maiti, I. B. (2013). An intergenic region shared by At4g35985 and At4g35987 in Arabidopsis thaliana is a tissue specific and stress inducible bidirectional promoter analyzed in transgenic arabidopsis and tobacco plants. PloS one, 8(11), e79622.

98. Bonke, M., Thitamadee, S., Mähönen, A. P., Hauser, M. T., & Helariutta, Y. (2003). APL regulates vascular tissue identity in Arabidopsis. Nature, 426(6963), 181–186.

99. Brady, S. M., Orlando, D. A., Lee, J. Y., Wang, J. Y., Koch, J., Dinneny, J. R., Mace, D., Ohler, U., & Benfey, P. N. (2007). A high-resolution root spatiotemporal map reveals dominant expression patterns. Science (New York, N.Y.), 318(5851), 801– 806.

100. Cayla, T., Le Hir, R., & Dinant, S. (2019). Live-Cell Imaging of Fluorescently Tagged Phloem Proteins with Confocal Microscopy. Methods in molecular biology (Clifton, N.J.), 2014, 95–108.

101. Chauvin, A., Caldelari, D., Wolfender, J. L., & Farmer, E. E. (2013). Four 13-lipoxygenases contribute to rapid jasmonate synthesis in wounded Arabidopsis thaliana leaves: a role for lipoxygenase 6 in responses to long-distance wound signals. The New phytologist, 197(2), 566–575.

102. Chen, L. Q., Qu, X. Q., Hou, B. H., Sosso, D., Osorio, S., Fernie, A. R., & Frommer, W. B. (2012). Sucrose efflux mediated by SWEET proteins as a key step for phloem transport. Science (New York, N.Y.), 335(6065), 207–211.

103. Clay, N. K., & Nelson, T. (2005). Arabidopsis thickvein mutation affects vein thickness and organ vascularization, and resides in a provascular cell-specific spermine synthase involved in vein definition and in polar auxin transport. Plant physiology, 138(2), 767–777.

104. Dinkeloo, K., Boyd, S., & Pilot, G. (2018). Update on amino acid transporter functions and on possible amino acid sensing mechanisms in plants. Seminars in cell & developmental biology, 74, 105–113.

105. Fisher, K., & Turner, S. (2007). PXY, a receptor-like kinase essential for maintaining polarity during plant vascular-tissue development. Current biology: CB, 17(12), 1061–1066.

106. Funk, V., Kositsup, B., Zhao, C., & Beers, E. P. (2002). The Arabidopsis xylem peptidase XCP1 is a tracheary element vacuolar protein that may be a papain ortholog. Plant physiology, 128(1), 84–94.

107. Gotor, C., Cejudo, F. J., Barroso, C., & Vega, J. M. (1997). Tissue-specific expression of ATCYS-3A, a gene encoding the cytosolic isoform of O-acetylserine(thiol)lyase in Arabidopsis. The Plant journal: for cell and molecular biology, 11(2), 347–352.

108. Huang, L., Shi, X., Wang, W., Ryu, K. H., & Schiefelbein, J. (2017). Diversification of Root Hair Development Genes in Vascular Plants. Plant physiology, 174(3), 1697– 1712.

109. Illouz-Eliaz, N., Lande, K., Yu, J., Jow, B., Swift, J., Lee, T., Nobori, T., Castanon, R.G., Nery, J.R. and Ecker, J.R., 2023. Drought Recovery Induced Immunity Confers Pathogen Resistance. bioRxiv, pp.2023–02.

110. Johnson, C. S., Kolevski, B., & Smyth, D. R. (2002). TRANSPARENT TESTA GLABRA2, a trichome and seed coat development gene of Arabidopsis, encodes a WRKY transcription factor. The Plant cell, 14(6), 1359–1375.

111. Kamiya, T., Borghi, M., Wang, P., Danku, J. M., Kalmbach, L., Hosmani, P. S., Naseer, S., Fujiwara, T., Geldner, N., & Salt, D. E. (2015). The MYB36 transcription factor orchestrates Casparian strip formation. Proceedings of the National Academy of Sciences of the United States of America, 112(33), 10533–10538.

112. Candela, H., Martínez-Laborda, A., & Micol, J. L. (1999). Venation pattern formation in Arabidopsis thaliana vegetative leaves. Developmental biology, 205(1), 205–216.

113. Kirik, V., Schnittger, A., Radchuk, V., Adler, K., Hülskamp, M., & Bäumlein, H. (2001). Ectopic expression of the Arabidopsis AtMYB23 gene induces differentiation of trichome cells. Developmental biology, 235(2), 366–377.

114. Kirik, V., Lee, M. M., Wester, K., Herrmann, U., Zheng, Z., Oppenheimer, D., Schiefelbein, J., & Hulskamp, M. (2005). Functional diversification of MYB23 and GL1 genes in trichome morphogenesis and initiation. Development (Cambridge, England), 132(7), 1477–1485.

115. Lee, J. Y., Colinas, J., Wang, J. Y., Mace, D., Ohler, U., & Benfey, P. N. (2006). Transcriptional and posttranscriptional regulation of transcription factor expression in Arabidopsis roots. Proceedings of the National Academy of Sciences of the United States of America, 103(15), 6055–6060.

116. Liberman, L. M., Sparks, E. E., Moreno-Risueno, M. A., Petricka, J. J., & Benfey, P. N. (2015). MYB36 regulates the transition from proliferation to differentiation in the Arabidopsis root. Proceedings of the National Academy of Sciences of the United States of America, 112(39), 12099–12104.

117. Menand, B., Yi, K., Jouannic, S., Hoffmann, L., Ryan, E., Linstead, P., Schaefer, D. G., & Dolan, L. (2007). An ancient mechanism controls the development of cells with a rooting function in land plants. Science (New York, N.Y.), 316(5830), 1477–1480.

118. Mizukami, Y., & Ma, H. (1997). Determination of Arabidopsis floral meristem identity by AGAMOUS. The Plant cell, 9(3), 393–408.

119. Mølhøj, M., Jørgensen, B., Ulvskov, P., & Borkhardt, B. (2001). Two Arabidopsis thaliana genes, KOR2 and KOR3, which encode membrane-anchored endo-1,4-beta-D-glucanases, are differentially expressed in developing leaf trichomes and their support cells. Plant molecular biology, 46(3), 263–275.

120. Muñiz, L., Minguet, E. G., Singh, S. K., Pesquet, E., Vera-Sirera, F., Moreau-Courtois, C. L., Carbonell, J., Blázquez, M. A., & Tuominen, H. (2008). ACAULIS5 controls Arabidopsis xylem specification through the prevention of premature cell death. Development (Cambridge, England), 135(15), 2573–2582.

121. Nguyen, C. T., Kurenda, A., Stolz, S., Chételat, A., & Farmer, E. E. (2018). Identification of cell populations necessary for leaf-to-leaf electrical signaling in a wounded plant. Proceedings of the National Academy of Sciences of the United States of America, 115(40), 10178–10183.

122. Oppenheimer, D. G., Pollock, M. A., Vacik, J., Szymanski, D. B., Ericson, B., Feldmann, K., & Marks, M. D. (1997). Essential role of a kinesin-like protein in Arabidopsis trichome morphogenesis. Proceedings of the National Academy of Sciences of the United States of America, 94(12), 6261–6266.

123. Pedersen, D. S., Coppens, F., Ma, L., Antosch, M., Marktl, B., Merkle, T., Beemster, G. T., Houben, A., & Grasser, K. D. (2011). The plant-specific family of DNA-binding proteins containing three HMG-box domains interacts with mitotic and meiotic chromosomes. The New phytologist, 192(3), 577–589.

124. Pilot, G., Pratelli, R., Gaymard, F., Meyer, Y., & Sentenac, H. (2003). Five-group distribution of the Shaker-like K+ channel family in higher plants. Journal of molecular evolution, 56(4), 418–434.

125. Redovniković, I. R., Textor, S., Lisnić, B., & Gershenzon, J. (2012). Expression pattern of the glucosinolate side chain biosynthetic genes MAM1 and MAM3 of Arabidopsis thaliana in different organs and developmental stages. Plant physiology and biochemistry: PPB, 53, 77–83.

126. Sasabe, M., Ishibashi, N., Haruta, T., Minami, A., Kurihara, D., Higashiyama, T., Nishihama, R., Ito, M., & Machida, Y. (2015). The carboxyl-terminal tail of the stalk of Arabidopsis NACK1/HINKEL kinesin is required for its localization to the cell plate formation site. Journal of plant research, 128(2), 327–336.

127. Sawa, S., Watanabe, K., Goto, K., Liu, Y. G., Shibata, D., Kanaya, E., Morita, E. H., & Okada, K. (1999). FILAMENTOUS FLOWER, a meristem and organ identity gene of Arabidopsis, encodes a protein with a zinc finger and HMG-related domains. Genes & development, 13(9), 1079–1088.

128. Sawchuk, M. G., Donner, T. J., Head, P., & Scarpella, E. (2008). Unique and overlapping expression patterns among members of photosynthesis-associated nuclear gene families in Arabidopsis. Plant physiology, 148(4), 1908–1924.

129. Scarpella, E., Francis, P., & Berleth, T. (2004). Stage-specific markers define early steps of procambium development in Arabidopsis leaves and correlate termination of vein formation with mesophyll differentiation. Development (Cambridge, England), 131(14), 3445–3455.

130. Schellmann, S., Schnittger, A., Kirik, V., Wada, T., Okada, K., Beermann, A., Thumfahrt, J., Jürgens, G., & Hülskamp, M. (2002). TRIPTYCHON and CAPRICE mediate lateral inhibition during trichome and root hair patterning in Arabidopsis. The EMBO journal, 21(19), 5036–5046.

131. Schuster, J., Knill, T., Reichelt, M., Gershenzon, J., & Binder, S. (2006). Branched-chain aminotransferase4 is part of the chain elongation pathway in the biosynthesis of methionine-derived glucosinolates in Arabidopsis. The Plant cell, 18(10), 2664– 2679.

132. Stacey, M. G., Osawa, H., Patel, A., Gassmann, W., & Stacey, G. (2006). Expression analyses of Arabidopsis oligopeptide transporters during seed germination, vegetative growth and reproduction. Planta, 223(2), 291–305.

133. Stadler, R., & Sauer, N. (2019). The AtSUC2 Promoter: A Powerful Tool to Study Phloem Physiology and Development. Methods in molecular biology (Clifton, N.J.), 2014, 267–287.

134. Susek, R. E., Ausubel, F. M., & Chory, J. (1993). Signal transduction mutants of Arabidopsis uncouple nuclear CAB and RBCS gene expression from chloroplast development. Cell, 74(5), 787–799.

135. Lu, P., Porat, R., Nadeau, J. A., & O’Neill, S. D. (1996). Identification of a meristem L1 layer-specific gene in Arabidopsis that is expressed during embryonic pattern formation and defines a new class of homeobox genes. The Plant cell, 8(12), 2155– 2168.

136. Sessions, A., Weigel, D., & Yanofsky, M. F. (1999). The Arabidopsis thaliana MERISTEM LAYER 1 promoter specifies epidermal expression in meristems and young primordia. The Plant journal: for cell and molecular biology, 20(2), 259–263.

137. Vanzin, G. F., Madson, M., Carpita, N. C., Raikhel, N. V., Keegstra, K., & Reiter, W. D. (2002). The mur2 mutant of Arabidopsis thaliana lacks fucosylated xyloglucan because of a lesion in fucosyltransferase AtFUT1. Proceedings of the National Academy of Sciences of the United States of America, 99(5), 3340–3345.

138. Wendrich, J. R., Yang, B., Vandamme, N., Verstaen, K., Smet, W., Van de Velde, C., Minne, M., Wybouw, B., Mor, E., Arents, H. E., Nolf, J., Van Duyse, J., Van Isterdael, G., Maere, S., Saeys, Y., & De Rybel, B. (2020). Vascular transcription factors guide plant epidermal responses to limiting phosphate conditions. Science (New York, N.Y.), 370(6518), eaay4970.

139. Wenzel, C. L., Schuetz, M., Yu, Q., & Mattsson, J. (2007). Dynamics of MONOPTEROS and PIN-FORMED1 expression during leaf vein pattern formation in Arabidopsis thaliana. The Plant journal: for cell and molecular biology, 49(3), 387– 398.

140. Xu, W., Purugganan, M. M., Polisensky, D. H., Antosiewicz, D. M., Fry, S. C., & Braam, J. (1995). Arabidopsis TCH4, regulated by hormones and the environment, encodes a xyloglucan endotransglycosylase. The Plant cell, 7(10), 1555–1567.

141. Yanofsky, M. F., Ma, H., Bowman, J. L., Drews, G. N., Feldmann, K. A., & Meyerowitz, E. M. (1990). The protein encoded by the Arabidopsis homeotic gene agamous resembles transcription factors. Nature, 346(6279), 35–39.

142. Zheng, H., Rowland, O., & Kunst, L. (2005). Disruptions of the Arabidopsis Enoyl-CoA reductase gene reveal an essential role for very-long-chain fatty acid synthesis in cell expansion during plant morphogenesis. The Plant cell, 17(5), 1467–1481

